# Systematic modeling of phenotypic drug response profiles in patient-derived organoids

**DOI:** 10.64898/2026.07.29.741618

**Authors:** Seungil Kim, Einar Bjarki Gunnarsson, Michael E. Doche, Yuyuan Zhou, Yu-Kai Huang, Scott Valena, Pratiksha Kshetri, Elizabeth Elton, Nolan Ung, Abigail Coleman, Benedikt Vilji Magnússon, Niki Torab, Lara S. Papasian, Brandon Choi, Eileen Fung, Hanaa Knaneh-Monem, Jasmine Foo, Shannon M. Mumenthaler

**Affiliations:** Ellison Medical Institute, Los Angeles, CA. USA; Applied Mathematics Division, Science Institute, University of Iceland, Reykjavík, Iceland; School of Mathematics, University of Minnesota, Twin Cities, MN. USA; Department of Psychology, College of Letters and Sciences, University of California, Los Angeles, CA. USA; Keck School of Medicine, University of Southern California, Los Angeles, CA. USA; Department of Biomedical Engineering, Viterbi School of Engineering, University of Southern California, Los Angeles, CA. USA

## Abstract

Patient-derived tumor organoids provide a physiologically relevant 3D disease model for preclinical drug discovery, surpassing the limitations of conventional 2D cell lines. To better capture the dynamic nature of organoid drug responses, we developed a new systematic evaluation method called SCOPE (Systematic Classification of Organoids for Phenotypic Evaluation), harnessing phenotypic assessments from multi-timepoint 3D imaging data. By integrating artificial intelligence (AI)-based image analysis of organoid viability with tracking and mathematical modeling of organoid growth over time, we captured temporal-and dose-dependent dynamics of phenotypic changes, culminating in two novel metrics: a combined growth and viability (*GV*) score as well as a cytostatic-cytotoxic transition range (CCTR) that separates drug effects on organoid growth and viability. Our approach supports classification of specific drug responses into four distinct phenotypic groups: (1) cytotoxic, (2) cytostatic plus cytotoxic, (3) late cytotoxic, and (4) cytostatic. This novel drug evaluation system can identify previously unknown drug effects or new therapeutic use cases for existing drugs, facilitating the design of alternative therapeutic options to overcome efficacy or drug resistance challenges and improving the clinical applicability of organoid-based drug discovery results.

## INTRODUCTION

Traditional drug discovery has relied on static cell viability analysis of two-dimensional (2D) cells treated with drugs. Although this approach allows for fast, high-throughput screening of large numbers of drug candidates, it lacks both temporal and spatial contexts that can provide additional information on the mechanisms of action and potential side effects ^1^. To overcome the limitations of 2D cell lines in capturing key aspects of human biology, three-dimensional (3D) *in vitro* models, such as spheroids, organoids and organs-on-chips, have been developed to more accurately recapitulate cell-cell interactions, tissue morphology, and the complex microenvironment. These 3D culture systems are increasingly utilized to evaluate the efficacy and toxicity of new drug candidates ^2,3^. In the oncology setting, patient-derived organoids (PDOs) hold significant promise for drug studies, as they can mirror clinical responses at the individual patient level and help predict treatment outcomes ^4,5^.

Conventional chemotherapies targeting tumor cells, which are predominantly cytotoxic, have historically been the cornerstone of treatment. In contrast, targeted therapies act largely through cytostatic mechanisms, by interfering with cell signaling pathways to inhibit tumor cell proliferation without directly inducing cytotoxicity. These distinct mechanisms of action give rise to fundamentally different tumor cell population dynamics, which must be explicitly considered when evaluating drug responses and optimizing therapeutic regimens. For example, by reducing non-specific cytotoxicity, cytostatic therapies may have lower side effects; however, because they suppress cell cycle progression rather than increase cell death, longer term treatments may be necessary for achieving tumor reduction, in turn increasing the risk of developing drug-resistant subpopulations ^6^. To capture these nuances, more physiologically relevant models and dynamic readouts are needed to assess therapeutic responses and predict clinical outcomes across different drug modalities and targets.

Advances in imaging technology have transformed high-content phenotypic screening into a powerful drug discovery tool, offering unprecedented resolution across spatial and temporal dimensions. Recently, AI-based approaches have expedited 3D image analysis processes with pre-trained neural networks (NNs) capable of identifying cells automatically and extracting rich cellular and morphological measurements from large-scale datasets ^7,8^. Coupling high-resolution imaging with AI-driven analysis and mathematical modeling can provide a powerful approach to go beyond snapshots of complex and evolving biological systems. Mathematical modeling offers a principled framework for integrating longitudinal imaging data with mechanistic understanding of the underlying biological processes, allowing us to reconstruct and interpret the continuous dynamics of organoid growth and drug response ^9,10^. We recently developed an integrated imaging and modeling framework to show that the Gompertz model can be used to describe tumor organoid growth and to quantify both intra-and inter-patient heterogeneities in the absence of treatment ^11^. Our results were consistent with previous evidence showing broad applicability of the Gompertz model for describing tumor growth across *in vitro, in vivo,* and clinical settings ^12–17^. Here, we extend this framework to develop new metrics that enable phenotypic classifications of dynamic drug responses of PDOs across different dose ranges.

By combining patient-derived 3D models with multi-scale imaging and phenotypic classification, here we establish a scalable platform for drug evaluation, referred to as **S**ystematic **C**lassification of **O**rganoids for **P**henotypic **E**valuation **(SCOPE),** that captures the temporal complexity of drug response and reveals patterns not detectable with static measurements. Our method enables patient and dose-specific assessments of functional drug response, providing richer insights into treatment design beyond molecular alterations.

## RESULTS

### Multi-modal imaging-based PDO evaluation system can provide comprehensive assessments of drug responses

Our new phenotypic evaluation method, SCOPE, uses dynamic 3D label-free imaging of PDOs. NN image analysis of multi-time and multi-dose data coupled with mathematical modeling stratifies patients based on specific drug response types **(Figure 1, Figure S1)**. This was demonstrated using a set of PDO lines from six different colorectal cancer (CRC) patient samples (EICL-000U7 (U7), EICL-000UA (UA), EICL-000UP (UP), EICL-000US (US), EICL-000UX (UX), and EICL-000V8 (V8)) **(Table S1)**. Initially, PDOs were treated with two standard chemotherapies that are routinely used clinically for CRC: SN38, the active metabolite of irinotecan, and 5-fluorouracil (5FU) **(Figure 2, Figure S2, S3A)**. For each PDO line and treatment condition, a previously described AI image analysis workflow ^11^ was used to collect multi-modal time series data describing the area, morphology and viability (live/dead) of individual organoids on Day 0 (pre-treatment) and Days 3 and 5 (post-treatment) **(Figure S1B-E, Video S1)**. **Figure 2A** shows representative images of SN38-treated PDOs comparing drug effects across two doses. The Gompertz model was applied to the time series data to estimate individual organoid growth rates, which were then used to define a *growth score* quantifying the drug effect on organoid growth for each PDO line and treatment condition. The growth score is based on model predictions of how large the organoids will be when they stop growing. A *viability score* was also defined using the proportion of organoids that were alive both on Days 3 and 5 according to NN analysis. We generated two distinct dose response curves (DRCs) using these metrics, separating the effects of the drug on organoid growth and viability. To compare these curves to a more conventional measure of drug effect on total cell viability, we also performed a separate luminescence assay using the CellTiter-Glo (CTG) reagent (**Figure 2B, S3B**). For each DRC, 50 % inhibition concentrations were computed, which we identify as IC_50_-growth, IC_50_-viability, and IC_50_-CTG.

**Figure 1:**
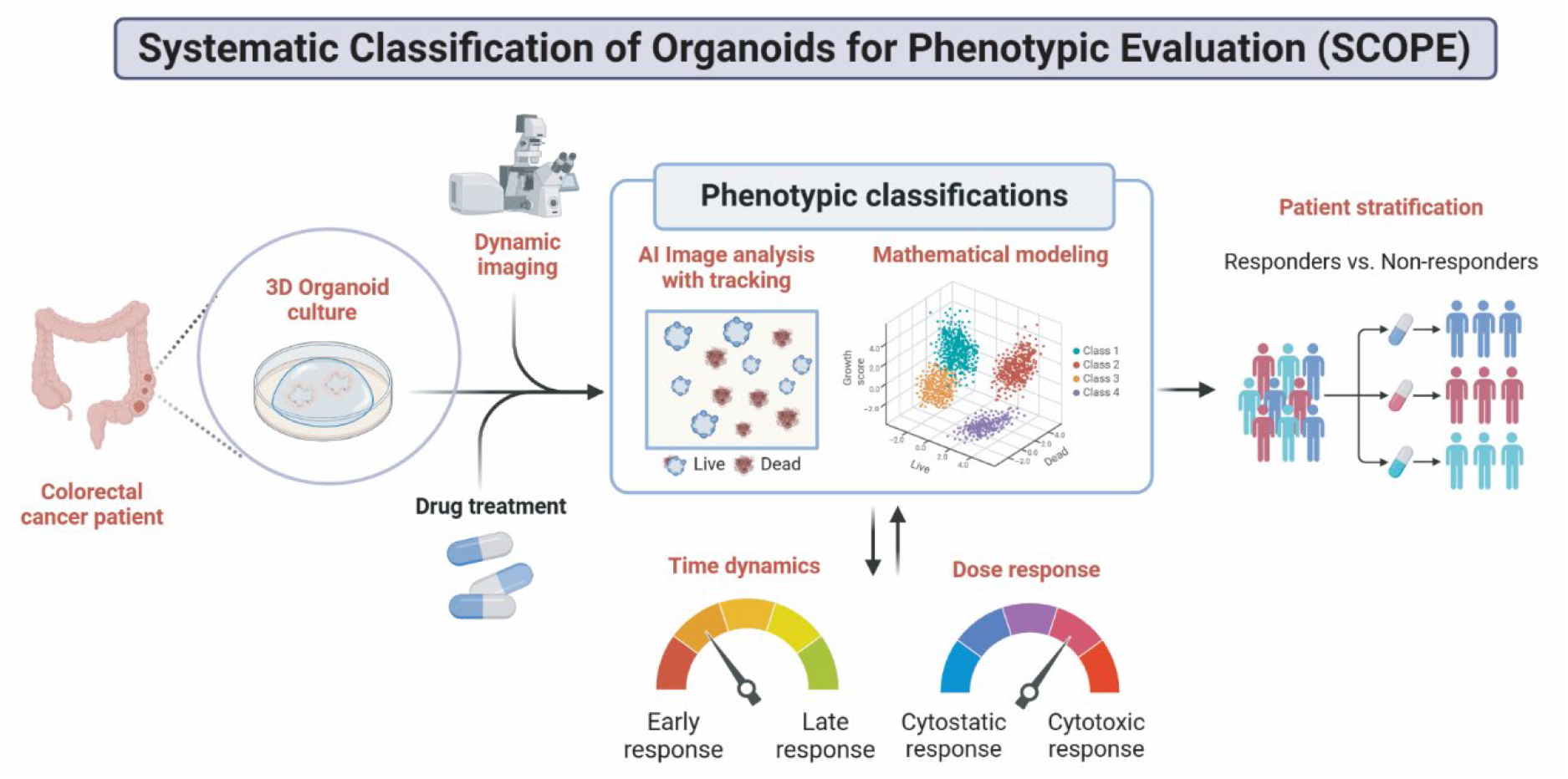
Phenotypic evaluation of PDOs with SCOPE. SCOPE integrates dynamic imaging, AI image analysis, and mathematical modeling to evaluate PDO drug responses with comprehensive information using increased spatial and temporal resolutions. Patient and dose-dependent phenotypic classifications can support responder stratification and inform treatment strategies. Created with BioRender software (https://BioRender.com)

**Figure 2:**
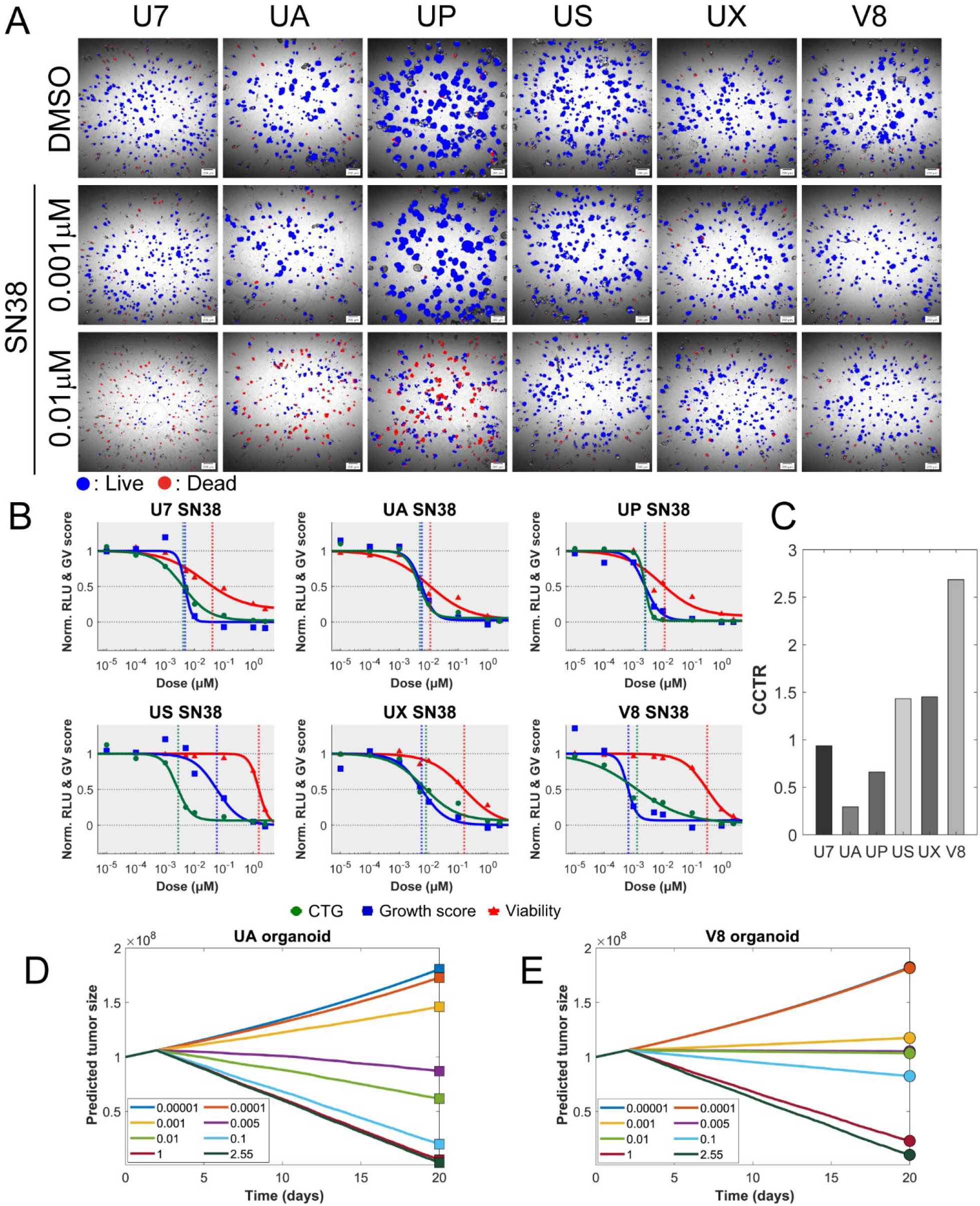
Capturing growth and viability metrics to assess drug response in PDOs. **A.** Representative NN-processed Day 5 images of six PDOs treated with SN38. Two doses are shown to compare the difference in drug response with the full dose panel captured in Fig. S2. NN masks - Blue: Live PDOs, Red: Dead PDOs. Scale bars, 200 µm. **B.** SN38 dose response curves of six PDOs based on CTG (Green), growth (Blue) and viability (Red) scores. Three graphs are overlayed indicating the IC_50_-CTG (vertical green dotted lines), IC_50_-growth (vertical blue dotted lines), IC_50_-viability (vertical red dotted lines) positions. **C.** Comparison of CCTR in six PDOs with SN38. The differences between log_10_ IC_50_-viability and log_10_ IC_50_-growth in six PDOs with SN38 are shown in the bar graph. CCTR = log_10_ (IC_50_ viability) – log_10_ (IC_50_ growth). **D, E.** Simulated UA patient tumor response and V8 patient tumor response to SN38 across experimentally tested doses, according to a simplified mathematical model relating PDO growth to tumor growth. CCTR: Cytostatic-cytotoxic transition range, CCTR = log_10_ (IC_50_ viability) – log_10_ (IC_50_ growth), Norm.: Normalized, RLU: Relative Luminescence Unit, GV: Growth-Viability.

The growth score DRC (blue curves) for SN38 closely resembled the CTG DRC (green curves) for all but the US PDO. However, the degree of separation between the viability score DRC (red curves) and the growth score DRC varied between patients (**Figure 2B**). We quantified this separation by computing the distance between the IC_50_-growth and IC_50_-viability doses **(Data S1)** for each PDO line, a metric we refer to as the *cytostatic-cytotoxic transition range* (CCTR) **(Figure 2C)**. A small CCTR indicates that IC_50_-growth and viability doses are close, and thus that organoid killing effects largely coincide with growth inhibition effects. A large CCTR indicates that growth inhibition can be achieved at low doses, whereas much larger doses are required to induce organoid death. For example, the response of UA PDOs to 0.01 µM of SN38 was a combination of partial growth inhibition and partial organoid killing, whereas the V8 PDO response was attributed to strong growth inhibition with little to no organoid killing (bottom row of **Figure 2A**). This difference in drug response, which is not apparent from the CTG DRC, is clearly indicated by a large difference in CCTR between UA and V8 PDOs **(Figure 2B-C)**. Individual organoid-level analysis of viability and growth rates confirmed that SN38 reduced organoid size and growth rates and increased dead classifications in a dose-dependent and PDO-specific manner **(Figure S4)**. In contrast to SN38, 5FU inhibited PDO growth with high doses but did not kill organoids effectively, representing typical characteristics of cytostatic drugs **(Figure S3)**. The IC_50_-growth doses varied across PDOs between the patients but were overlapping or close to the IC_50_ values from CTG except for the UX PDO. UX PDOs showed more live organoids under the highest dose, 100 µM, compared to the other PDO lines **(Figure S3B, Data S1)**. 5FU killed around half of the UP PDOs at the highest dose, but otherwise had limited effect on organoid viability, indicating large to infinite IC_50_-viability values **(Figure S3B).**

To illustrate the potential clinical relevance of the CCTR, we developed a simplified mathematical model of tumor growth based on clonal evolution ^18^, where individual organoids correspond to individual subclones in the tumor. In the model, the drug is assumed to affect the tumor by killing a certain proportion of the subclones and reducing the growth rates of the remaining subclones by a certain percentage. In this context, the IC_50_-growth dose can be interpreted as the dose at which subclone growth is inhibited by 50 %, and the IC_50_-viability dose is the dose at which half the tumor subclones are killed. We used this model to simulate the drug responses of the UA and V8 patient tumors to SN38. For the simulated UA tumor, the drug gradually affects tumor evolution with increasing doses, with tumor reduction first observed at 0.005 µM **(Figure 2D)**, which is the same dose showing a significant increase in dead organoids **(Figure S2, Figure S4)**. For the simulated V8 tumor, a sharp transition occurs between no effect at 0.0001 µM and almost complete growth inhibition at 0.001 µM, whereas the dose must be increased 1000-fold to 1 µM to achieve significant tumor reduction **(Figure 2E)** recapitulating the results from individual organoid analysis **(Figure S4B)**. This is due to the large CCTR for V8 **(Figure 2C)**. From a clinical perspective, a large CCTR can indicate that complete growth inhibition is possible using low non-toxic doses, whereas depending on the drug’s toxicity profile, alternative treatments or combinations with cytotoxic drugs must be explored to achieve significant tumor reduction for the patient in question. More broadly, patient-specific CCTR analysis can be linked to patient information and molecular characterization data to explain differences in clinical outcomes under a certain drug.

### Phenotypic classifications of PDOs can be used as a drug screening tool to evaluate drug responses

We next used our image analysis workflow to conduct a large-scale drug screen for a specific colorectal cancer PDO line, UP. The UP PDOs were treated with 166 FDA-approved anti-cancer compounds at a single dose (10 µM) and multi-modal time series data was collected as before. We combined the time series data with mathematical modeling to develop a three-dimensional visualization framework for dynamic drug response, called a *growth-viability* (*GV*) plot **(Figure 3A)**. The *GV* plot indicates for each drug the proportion of organoids that remain alive on Day 5 (*x*-coordinate), the proportion of organoids that are alive on Day 3 but die by Day 5 (*y*-coordinate), and organoid growth score with specific drugs based on the Gompertz model (*z*-coordinate). The proportion of organoids that have already died by Day 3 is implicit in the *GV* plot, since it can be obtained by subtracting the *x*-and *y*-coordinates from 1. Thus, the *GV* plot separates the effect of each drug on growth and viability while also indicating its temporal response dynamics. The controls, 0.1 % dimethyl sulfoxide vehicle (DMSO, blue triangle) and staurosporine (STA, pan-kinase inhibitor, inverted red triangle) appear in opposite corners of the *GV* plot. The position of DMSO indicates that all organoids remain alive on Day 5 with no growth inhibition, whereas the position of STA indicates that the drug has already killed the organoids by Day 3 without allowing any growth. 5FU is positioned almost directly below DMSO, indicating growth inhibition without substantial organoid killing (cytostatic effect), while SN38 is positioned close to STA, indicating a predominantly cytotoxic effect.

**Figure 3:**
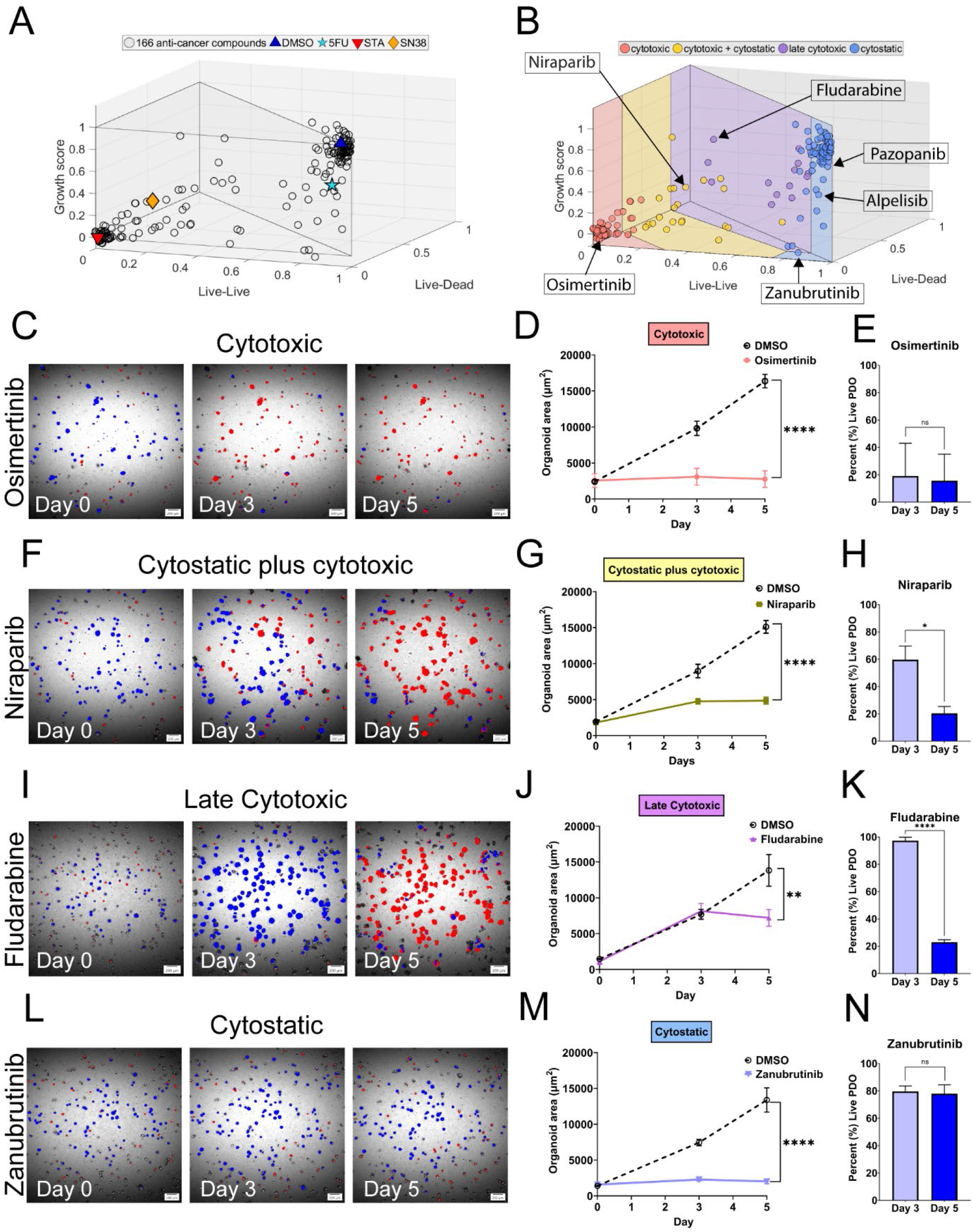
Phenotypic classifications of PDO drug responses from a drug screen. **A.** 3D *GV* plot of 166 drugs with controls (DMSO, 5FU, STA, SN38). **B.** 3D *GV* plot with phenotypic classifications (cytotoxic, cytostatic plus cytotoxic, late cytotoxic, cytostatic). Example drugs of each class are indicated with arrows and annotations. **C.** Representative NN-processed images of osimertinib-treated PDOs. **D.** PDO area changes over time with osimertinib treatment. (DMSO Day 5: 16,366.49 µm^2^ ± 941.03 µm^2^, Osimertinib Day 5: 2,769.53 µm^2^ ± 1,155.03 µm^2^, n = 3, *\*\*\*\*p<0.0001*). Mean ± SD. Two-way ANOVA test. **E.** Percent of live PDOs with osimertinib treatment at Days 3 and 5 (19 % ± 24.02 %, 15.67 % ± 19.35 %, n = 3, *^ns^p=0.5085*). Mean ± SD. Paired t-test. **F.** Representative NN-processed images of niraparib-treated PDOs. **G.** PDO area changes over time with niraparib treatment. (DMSO Day 5: 15,117.98 µm^2^ ± 882.62 µm^2^, Niraparib Day 5: 4,858.67 µm^2^ ± 491.18 µm^2^, n = 3, *\*\*\*\*p<0.0001*). Mean ± SD. Two-way ANOVA test. **H.** Percent of live PDOs with niraparib treatment at Days 3 and 5 (59.67 % ± 10.02 %, 20.33 % ± 5.03 %, n = 3, *\*p=0.0197*). Mean ± SD. Paired t-test. **I.** Representative NN-processed images of fludarabine-treated PDOs. **J.** PDO area changes over time with fludarabine treatment. (DMSO Day 5: 13,828.18 µm^2^ ± 2,209.16 µm^2^, Fludarabine Day 5: 7,201.79 µm^2^ ± 1,171.45 µm^2^, n = 3, *\*\*p=0.0019*). Mean ± SD. Two-way ANOVA test. **K.** Percent of live PDOs with fludarabine treatment at Days 3 and 5 (97.33 % ± 2.52 %, 23 % ± 1.73 %, n = 3, ****p<0.0001). Mean ± SD. Paired t-test. **L.** Representative NN-processed images of zanubrutinib-treated PDOs. **M.** PDO area changes over time with zanubrutinib treatment. (DMSO Day 5: 13,379.99 µm^2^ ± 1,700.04 µm^2^, Zanubrutinib Day 5: 2,010.26 µm^2^ ± 306.69 µm^2^, n = 3, *\*\*\*\*p<0.0001*). Mean ± SD. Two-way ANOVA test. **N.** Percent of live PDOs with zanubrutinib treatment at Days 3 and 5 (79.67 % ± 4.04 %, 78 % ± 6.56 %, n =3, *^ns^p=0.7284*). Scale bars, 200 µm.

Across the 166 anti-cancer drugs, the *GV* plot revealed significant heterogeneity in dynamic drug response. We used the *GV* plot to identify and define four phenotypic drug response classes: (1) cytotoxic (light red), (2) cytostatic plus cytotoxic (light yellow), (3) late cytotoxic (light purple) and (4) cytostatic (light blue) (**Figure 3B**). Cytotoxic drugs (example: osimertinib) showed strong PDO killing effects already on Day 3 after drug application. For example, under osimertinib, PDOs did not increase in size after Day 0 (**Figure 3C, D**), and the percentage of live PDOs was low both on Days 3 and 5, suggesting that most PDOs were killed rapidly (**Figure 3E**). Cytostatic plus cytotoxic drugs (example: niraparib) showed elements of both growth inhibition and cytotoxicity, with a mix of live and dead organoids on Day 3 and 5 (**Figure 3F, G**). Under niraparib, PDO growth stopped after Day 3, and there was a significant reduction in live PDOs between Days 3 and 5 (**Figure 3H**). Late cytotoxic drugs (example: fludarabine) neither displayed early cytotoxic effects nor strong growth inhibition. Under fludarabine, initial PDO growth was similar to DMSO control, and few PDOs had been killed by Day 3. However, dramatic viability changes occurred between Day 3 and 5, killing most PDOs (**Figure 3I-K**). Cytostatic drugs (example: zanubrutinib) inhibited PDO growth without affecting viability. Zanubrutinib stopped PDO growth completely (**Figure 3L, M**), resulting in a growth score close to 0, without inducing PDO death (**Figure 3N**). All tested drugs are listed in **Data S2** with coordinates of each 3D summary vector for UP PDOs.

To qualitatively validate the phenotypic classification from our SCOPE method, we conducted a manual review of images and classified the drugs based on visual assessments (**Data S3**). This resulted in only four mismatches in classification out of 123 selected drugs (the remaining 43 drugs were deemed as having no effect by CTG and NN analysis), which included edge cases where a drug could reasonably be assigned to two different classes. In general, while the four drug response classes captured high-level variations in phenotypic response, the *GV* plot also revealed significant variations in response within each phenotypic class, as described in **Figure S5** and the **Supplemental text**. For example, the *GV* plot revealed a wide range of cytostatic effects (**Figure 3B**), which motivated us to divide cytostatic drugs into 43different groups based on their growth score: strong cytostatic effect (example: zanubrutinib), medium effect (example: alpelisib), and weak or no effect (example: pazopanib) (**Figure 3L-N, S5J-L)**.

We then investigated how the four different phenotypic classes related to drug target signaling pathways and known biological processes **(Data S3, Figure 4)**. Both the cytotoxic and late cytotoxic classes contain many DNA synthesis inhibitors that are used in the clinic as cytotoxic chemotherapy agents **(Figure 4A, C)**. The cytostatic plus cytotoxic class includes cytotoxic chemotherapy as well as kinase or cell cycle inhibitors. The fraction of kinase and cell cycle inhibitors is large compared to the cytotoxic class, suggesting that these inhibitors targeting cell proliferation pathways can induce cytostatic effects **(Figure 4B)**. Consistently, the cytostatic class has more kinase and cell cycle inhibitors, showing that cytostatic effects can be achieved by using these types of drugs without inducing a strong cytotoxic effect **(Figure 4D)**. The effects of cytostatic drugs vary depending on the targets. For example, ibrutinib targeting Bruton tyrosine kinase induced a strong growth inhibition in UP PDOs **(Figure 4E)**. Alpelisib (inhibitor of PI3 kinase) and ARRY-380 (inhibitor of HER2) showed medium growth inhibition effects **(Figure 4F, G)**. Tazemetostat (inhibitor of EZH2) displayed a weak growth inhibition **(Figure 4H).** indicating that SCOPE can also classify phenotypic drug responses by known target mechanisms. The observed correlation between phenotypic classification derived from SCOPE and known drug mechanisms suggests that organoid-based SCOPE analysis can be used to prioritize drug targets through unbiased screening approaches.

**Figure 4:**
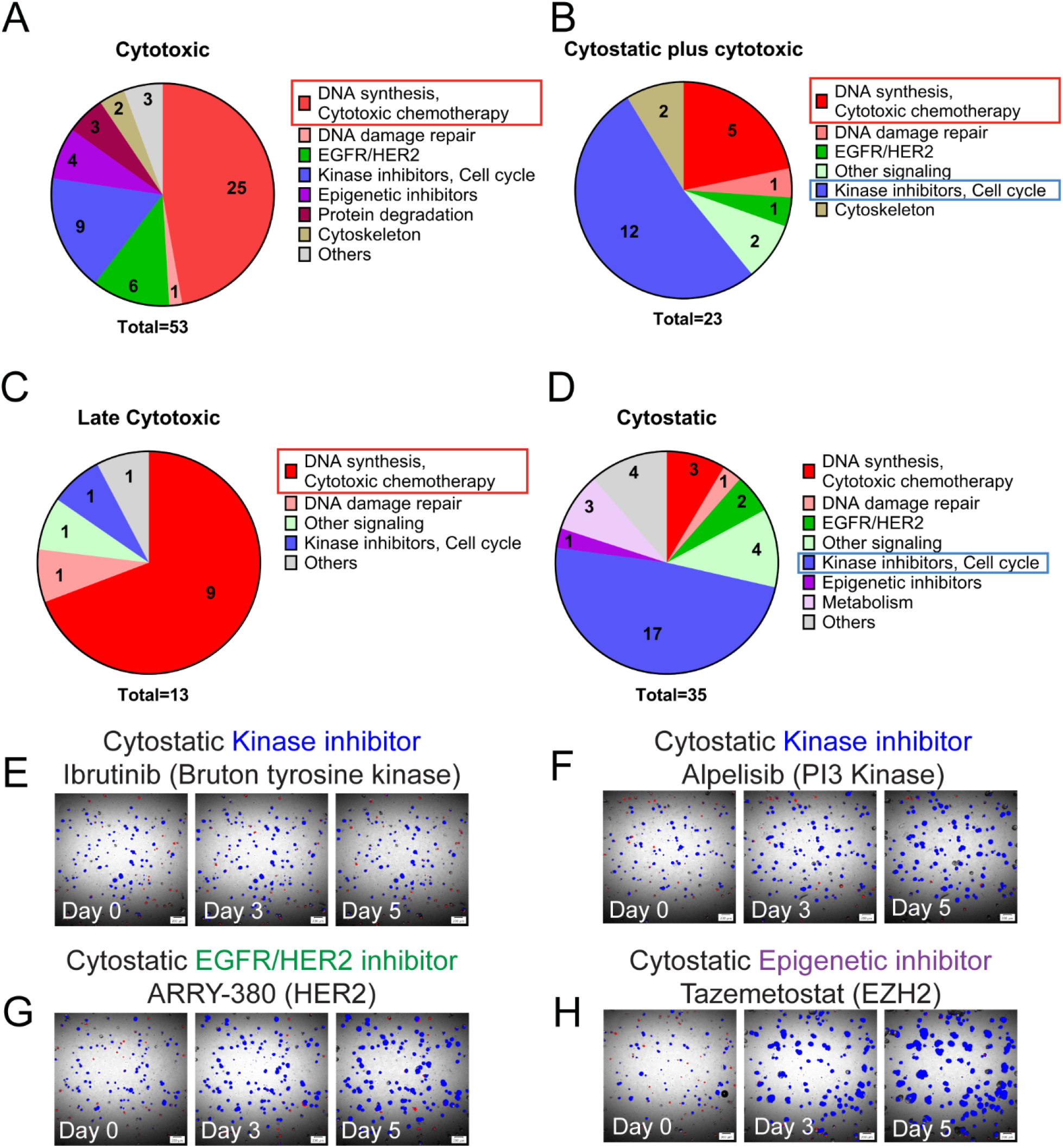
Phenotypic classifications of PDOs correlate with specific drug targets and related biological processes. **A.** Cytotoxic drugs are grouped based on the drug types and mechanisms. 53 drugs are divided in a pie chart with percentages according to the drug information. The number inside the chart shows the actual number of drugs. Drug types or pathways corresponding to the majority of drugs are indicated by a colored box. DNA synthesis inhibitors and cytotoxic chemotherapy agents are highlighted with a red box. **B.** Classification of cytostatic plus cytotoxic drugs based on the target pathways. Two major portions are highlighted by red (Cytotoxic chemotherapy, DNA synthesis inhibitors) and blue (cell cycle and kinase inhibitors) boxes. Total 23 drugs. **C.** Classification of late cytotoxic drugs. The major portion is highlighted with a solid red box (cytotoxic chemotherapy). Total 13 drugs. **D.** Classification of cytostatic drugs. The major drug types are highlighted with a blue box (cell cycle and kinase inhibitors). Total 35 drugs. **E.** Day 0, 3 and 5 NN-processed images of UP PDOs treated with ibrutinib (Bruton kinase inhibitor-treated), **F.** alpelisib (PI3 kinase inhibitor), **G.** ARRY-380 (HER2 inhibitor), and **H.** tazemetostat (EZH2 inhibitor). Scale bars, 200 µm.

### SCOPE distinguishes differential drug responses with same molecular targets

Although anti-cancer drugs target specific biological processes or molecules, the phenotypic outcomes can vary even between drugs targeting the same molecule. Epidermal growth factor receptor (EGFR) signaling is known as a critical pathway in multiple cancer types, including CRC. Although monoclonal antibody inhibitors of EGFR, such as cetuximab and panitumumab, are utilized in CRC treatment, acquired resistance remains a major challenge ^19,20^. Gefitinib and osimertinib, primarily developed for non-small cell lung cancer, target the EGFR tyrosine kinase domain to prevent downstream signaling pathways including RAS, RAF, MEK and ERK that regulate cell proliferation, differentiation and survival ^21^. Given the need for effective second-line therapies in CRC, exploring EGFR tyrosine kinase inhibitors (TKIs), such as gefitinib and osimertinib, that are available in the FDA-approved drug panel, offers a promising avenue for overcoming resistance and expanding the space of therapeutic options in CRC. Therefore, we selected these drugs to compare how SCOPE can be used to analyze differential responses to drugs with the same molecular targets, based on patient-specific phenotypic analysis.

Gefitinib and osimertinib treatments were applied to all six PDO lines. First, *GV* plots were generated to reveal dose-dependent phenotypic drug response trajectories **(Figure S6)** and to derive a dose-dependent phenotypic classification (**Figure 5A)**. Then, individual growth and viability DRCs were generated, and the IC_50_ values were computed as described before **(Figure 5B, Data S1)**. The separated DRCs showed that gefitinib induced strong growth inhibition in U7 (IC_50_-growth = 0.111 µM), UA (IC_50_-growth = 0.141 µM) and UP (IC_50_-growth = 0.320 µM) PDOs (**Figure 5B-first row**). Although the US PDOs (IC_50_-growth = 0.384 µM) were slightly more resistant to growth inhibition, their CCTR was small **(Figure 5C)**, indicating that cytotoxic effects coincided with growth inhibition. Gefitinib induced death in the US PDOs (IC_50_-viability = 0.514 µM) at a smaller dose than UP (IC_50_-viability = 3.091 µM), making the US PDOs more sensitive to the drug (**Figure 5B-second row left**). Both UX and V8 PDOs were clearly more resistant to gefitinib than the other PDOs, since viability was not affected even at the highest drug concentrations (**Figure 5B-second row middle and right, 5C**). The response dynamics and CCTRs for U7, UA and UP under osimertinib were similar to gefitinib **(Figure 5B, third row)**. However, contrary to gefitinib, the US PDOs (IC_50_-growth = 0.962 µM, IC_50_-viability = 7.387 µM) were more resistant to osimertinib than UP (IC_50_-growth = 0.328 µM, IC_50_-viability = 3.187 µM), both in terms of growth inhibition and cytotoxicity **(Figure 5B, forth row left)** with large CCTR values **(Figure 5D)**. UX and V8 were the most resistant to osimertinib, but we observed significant cytotoxic effects at the highest drug doses, suggesting that this drug can kill these gefitinib-resistant PDOs more effectively.

**Figure 5:**
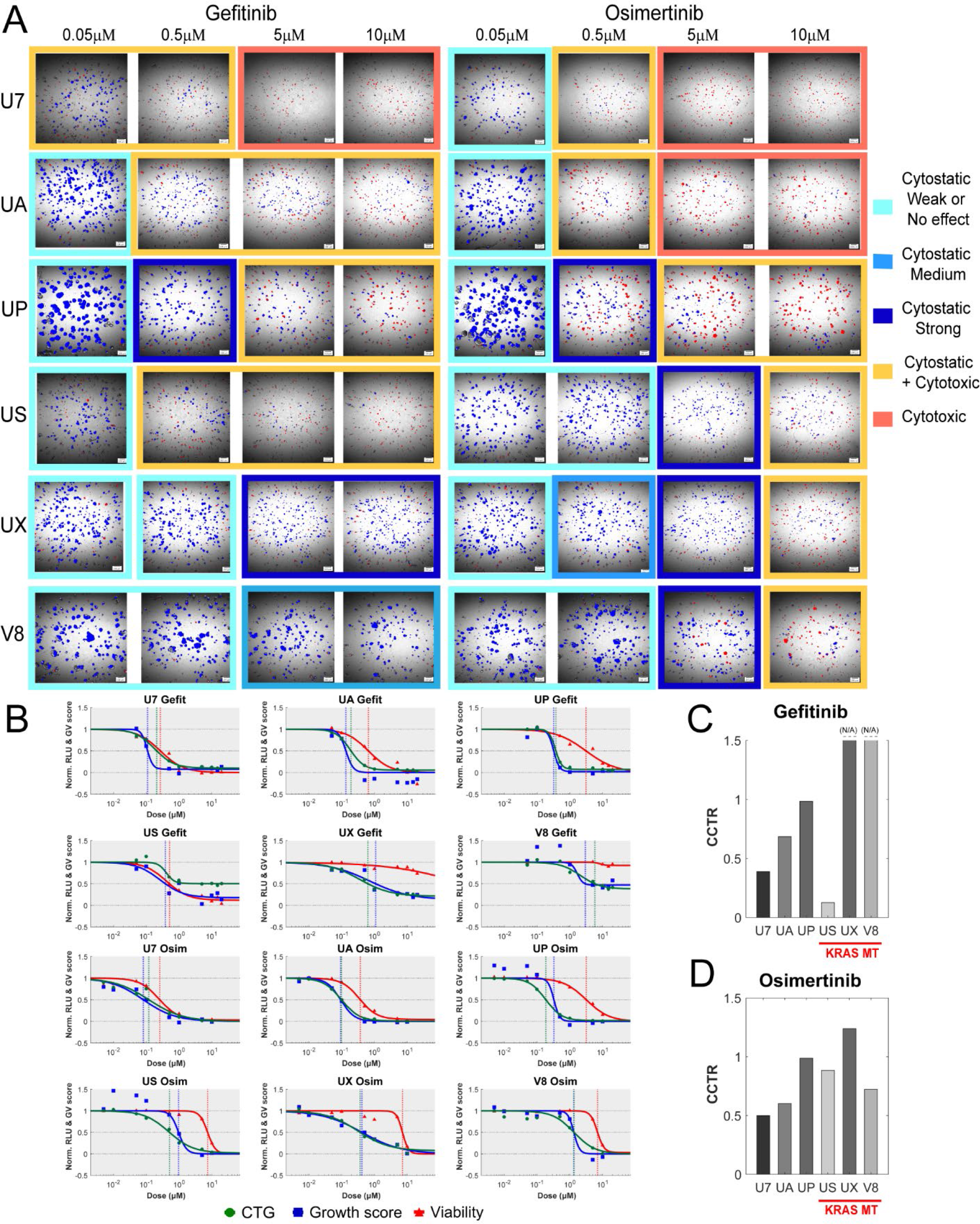
SCOPE analysis of gefitinib and osimertinib drug responses. **A.** Day 5 NN-processed images of six different PDOs treated with increasing doses of gefitinib and osimertinib. Each image is color-coded based on phenotypic classifications derived from 3D GV plots. Scale bars, 200 µm. **B.** DRCs of growth and viability scores as well as CTG of six PDOs treated with gefitinib and osimertinib. IC_50_-growth, IC_50-_viability and IC_50_-CTG are indicated with vertical blue, red and green dotted lines, respectively. **C.** CCTR of six PDOs with gefitinib. UX and V8 PDOs did not achieve 50% viability reduction at the highest tested dose, indicating large to infinite CCTRs. This is annotated by the horizontally dotted lines with (N/A - not available) sign. **D.** CCTR of six PDOs with osimertinib. KRAS mutant (MT) PDOs are underscored with red line.

For both drugs, the U7, UA and UP PDOs had lower IC_50_-growth values than US, UX and V8 PDOs **(Data S1)**. US PDOs showed a low (sensitive) IC_50_-growth (0.384 µM) with gefitinib similar to U7 (0.111 µM), UA (0.135 µM) and UP (0.320 µM) but the IC_50_-growth with osimertinib (0.962 µM) was higher (resistant) than UX PDOs (0.415 µM). UP, US, UX and V8 PDO lines generally had larger CCTRs than U7 and UA, excluding the US PDO for gefitinib **(Figure 5C**, **5D, Table S1)**. KRAS is one of the RAS family proteins and plays crucial roles in the EGFR signaling as a downstream mediator controlling cellular growth and proliferation ^22^. KRAS mutations are also known to be associated with refractory outcomes in patients treated with EGFR TKIs ^23,24^. KRAS mutations were not present in U7, UA and UP PDOs, classifying them as KRAS wild type. Within this group, the UP PDO stood out as having a higher IC_50_-growth and a larger CCTR. Even though a KRAS mutation was not identified in UP PDOs, there was an amplification of the KRAS gene **(Table S1)**. This difference in the copy number of KRAS (Copy number variation, CNV = 1.464) compared to the other KRAS wild type PDOs (U7, UA CNV = 0) potentially contributes to the higher level of resistance in UP PDOs. On the other hand, the sensitivity of US PDOs to gefitinib was noteworthy given the KRAS mutant status of the US PDO **(Table S1)**. In general, PDO groupings based on the IC_50_-growth doses and CCTRs correlate with their KRAS mutational status, with U7 and UA (KRAS wild type) being sensitive to both drugs and UP (KRAS amplification) as well as US, UX, V8 (KRAS mutants) being more resistant ; however there are exceptions, such as the US PDO with gefitinib, that highlight molecular alterations alone do not fully account for drug response **(Figure 5C**, **5D, Table S1)**.

## DISCUSSION

Significant challenges remain in overcoming the current high failure rate of candidate drugs in clinical trials. Addressing these challenges requires progress on multiple fronts: the development of more predictive and clinically relevant preclinical models for drug testing; more robust, reproducible and validated assays to measure phenotypic responses; integration of computational and AI approaches for the objective interpretation of assay outputs; and practical workflows that balance experimental complexity with the need for screening efficiency. PDOs are a promising model for addressing some of these challenges, as they better capture patient heterogeneity and tumor biology in phenotypic drug screens.

One area of focus is on establishing standard experimental procedures and quality control (QC) metrics to account for the complexities and heterogeneity of these patient-derived models ^25,26^. In our study, we established a novel automated phenotypic screening workflow using label-free 3D imaging to visually ensure consistency and reproducibility across culture plates and PDO lines. Coupling our approach with an endpoint viability assay enabled comparisons between phenotypic readouts and a traditional organoid-based viability assessment ^27^. We used pre-trained NNs and recorded macros for batch image analysis, thereby increasing efficiency as well as reducing human error and analysis time. Previously, Betge *et. al.* reported drug-induced phenotypic profiles in CRC organoids with over 500 small molecules ^28^. They also used an imaging-based method with a pre-trained random forest classifier. However, their training and feature detection relied on fluorescence staining with three different channels at a single timepoint after drug perturbation. In contrast, our SCOPE method analyzes three timepoints as a time series using a trained NN to track individual organoid changes and capture transitions in organoid growth, morphology, and viability without any fluorescence labeling.

PDOs from multiple cancer types have been shown to predict clinical responses to various drug modalities, including chemotherapy, radiation, and targeted therapies, using co-clinical trial approaches ^29–32^. A study by Tan et al. demonstrated a unified framework using PDOs to predict clinical responses to standard therapies in patients with metastatic CRC ^33^. They developed an imaging-based workflow to test drugs in PDOs with a rapid turn-around time. The PDO results were promising, demonstrating strong correlations to patient responses. Their method measured drug response by quantifying changes in mean PDO size over time as an indicator of viability. While this offers a single layer of information, we demonstrate with SCOPE that the ability to distinguish drug effects based on growth and viability (*GV* plot) provides additional insights into cytostatic and cytotoxic responses. We derived these features from integrating imaging-based phenotypic readouts and mathematical modeling-based interpretations. The *GV* plot from SCOPE can provide phenotypic classifications of multiple drugs in a single PDO, compare phenotypic responses to the same drug across PDOs, and group PDOs as responders or non-responders to specific drugs.

Another recent study by Yang *et al.* used mathematical modeling to compare the longitudinal response of pancreatic cancer PDOs to chemo-and radio-therapy, and to investigate the interplay between these two therapies ^34^. For chemotherapy, they applied the drug combination FOLFIRINOX at a single dose, and performed imaging on days 0, 2, 4, 6 and 7. They modeled organoid growth using the logistic growth model and assumed that the drug would reduce the growth rates of individual organoids according to the Norton-Simon hypothesis. In contrast, we performed both a large-scale drug screen with a single PDO line and multiple-dose experiments involving six PDOs treated with standard chemotherapy drugs (SN38, 5FU) and EGFR TKIs (gefitinib, osimertinib). In our analysis, we separated the drug effect on growth and viability, and we defined a growth score for each PDO line and treatment condition based on Gompertz growth modeling^34^. We generated distinct growth and viability DRCs, and introduced a novel metric, the CCTR, defined as the logarithmic difference between the IC_50_-viability and IC_50_-growth doses. The CCTR can provide mechanistic insights into drug selection and dose ranges needed for inducing cytostatic or cytotoxic effects. Interestingly, both Tan *et. al.* and Yang *et. al.* included combination therapy regimens in their PDO prediction studies, highlighting a potential avenue for the SCOPE method to explore novel drug pairs to improve patient outcomes ^33,34^. Dose-dependent phenotypic classifications from SCOPE can inform when higher doses or combination strategies using drugs of different classes may be needed.

Our high-throughput workflow produced data on growth and viability at three time points, which enabled us to visualize temporal drug response dynamics and identify drug phenotypes via the three-dimensional *GV* plot. For drug screening applications, increasing preclinical model complexity and spatiotemporal resolution can provide deeper insights and more precise estimates of longitudinal growth and viability effects; however, it can also increase experimental and analytical burdens and complicate visualization and categorization of drug responses. Our workflow is designed to flexibly balance workload with desired resolution and biological detail. We can: (1) incorporate traditional endpoint assays along with imaging and cell viability, (2) perform a two-step drug screening process, initially down selecting the number of drugs using an endpoint assay and then applying SCOPE, and (3) apply the full SCOPE profiling across all drugs. By integrating dynamic imaging with mathematical modeling, SCOPE reframes organoid-based drug screening from a single-outcome assay into a quantitative framework for evaluating dynamic drug responses across many cancer types.

Patient selection and stratification based on genetic information and molecular characterizations of biomarkers are used to guide clinicians in the design of optimal treatment plans with specific drugs. However, simple classifications as responders or non-responders based on this information may be neither accurate nor clinically relevant. Here we showed that SCOPE analysis involving IC_50_-growth and viability doses and CCTRs suggested patient groupings that broadly corresponded with KRAS mutational and amplification status, yet also revealed additional insights beyond those reflected by molecular alterations, thus providing complementary information for assessing drug responsiveness.

## ACKNOWLEDGEMENTS

We acknowledge C. Ambrose, R. Lau for PDO culture and CTG assay. K. Patsch and M. Purschke for Project discussion. E. Fong for review & editing. J. K. Schmid for help with data analysis. This study was supported by Innovative partnership in multi-scale bioimaging with Olympus-Evident. USC Dr. H.J. Lenz provided colorectal cancer patient tissues for generating PDOs. NCI Approved Oncology Drug plates were provided by the Developmental Therapeutics Program, Division of Cancer Treatment and Diagnosis, National Cancer Institute. The work of EBG was supported in part by NIH/NCI grant R01 CA241137.

## AUTHOR CONTRIBUTIONS

Conceptualization: S.K., E.B.G., J.F. and S.M.M., Methodology: S.K., E.B.G. and M.D., Investigation: S.K., E.B.G., M.D., Y.Z., Y.H., S.V., P.K. and B.C., Formal analysis: S.K., E.B.G., E.E., N.U., A.C., B.V.M., N.T. and L.S.P., Visualization: S.K., E.B.G., E.E., N.U., A.C., Software: E.B.G. and B.V.M., Data Curation: E.E. and N.U., Project administration: H.K.M., Supervision: J.F. and S.M.M., Writing—original draft: S.K. and E.B.G., Writing— review & editing: S.K., E.B.G., M.D., E.F., K.H., J..F and S.M.M.

## DECLARATIONS OF INTERESTS

Authors declare that they have no competing interests.

## STAR★METHODS

## KEY RESOURCES TABLE

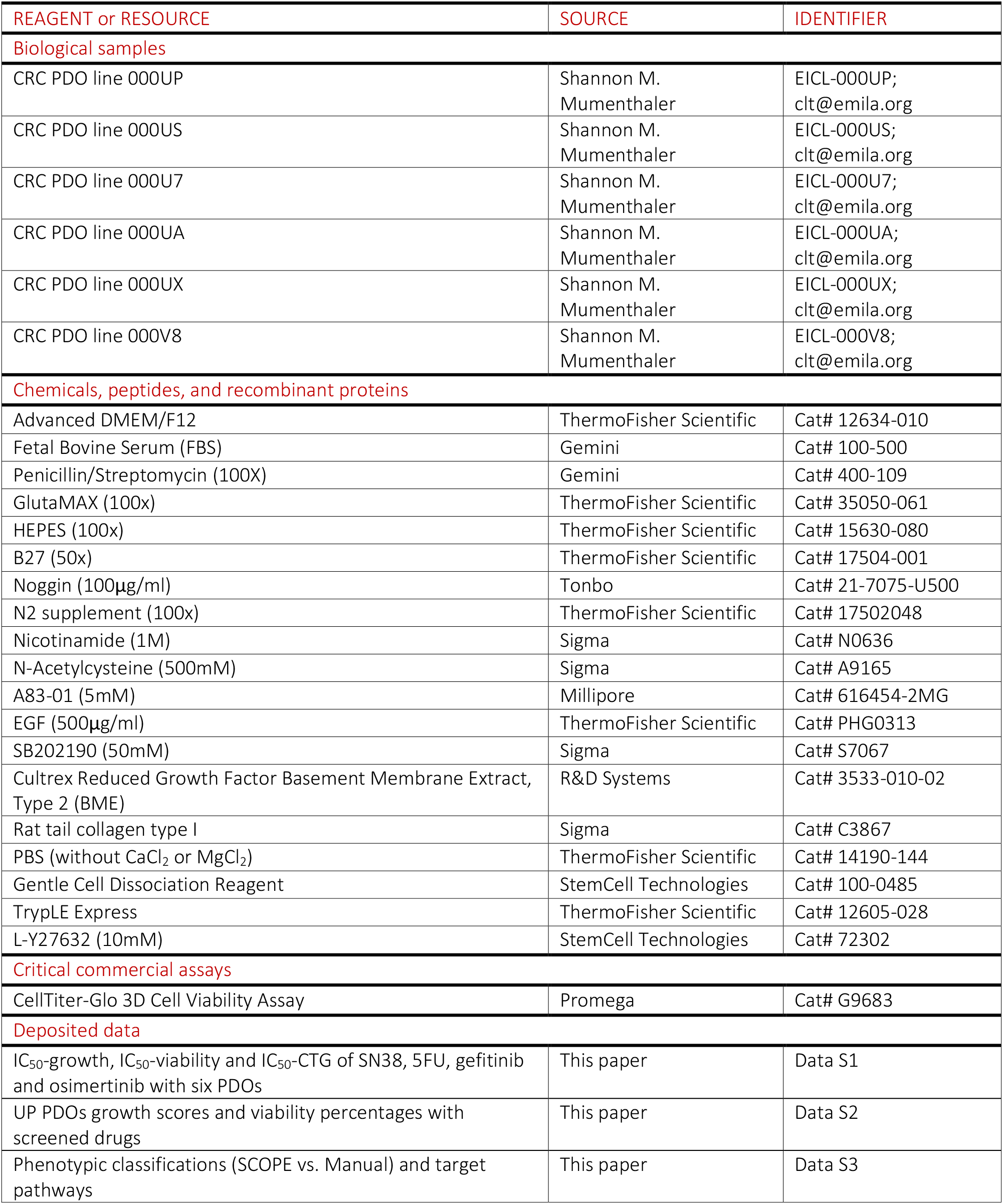
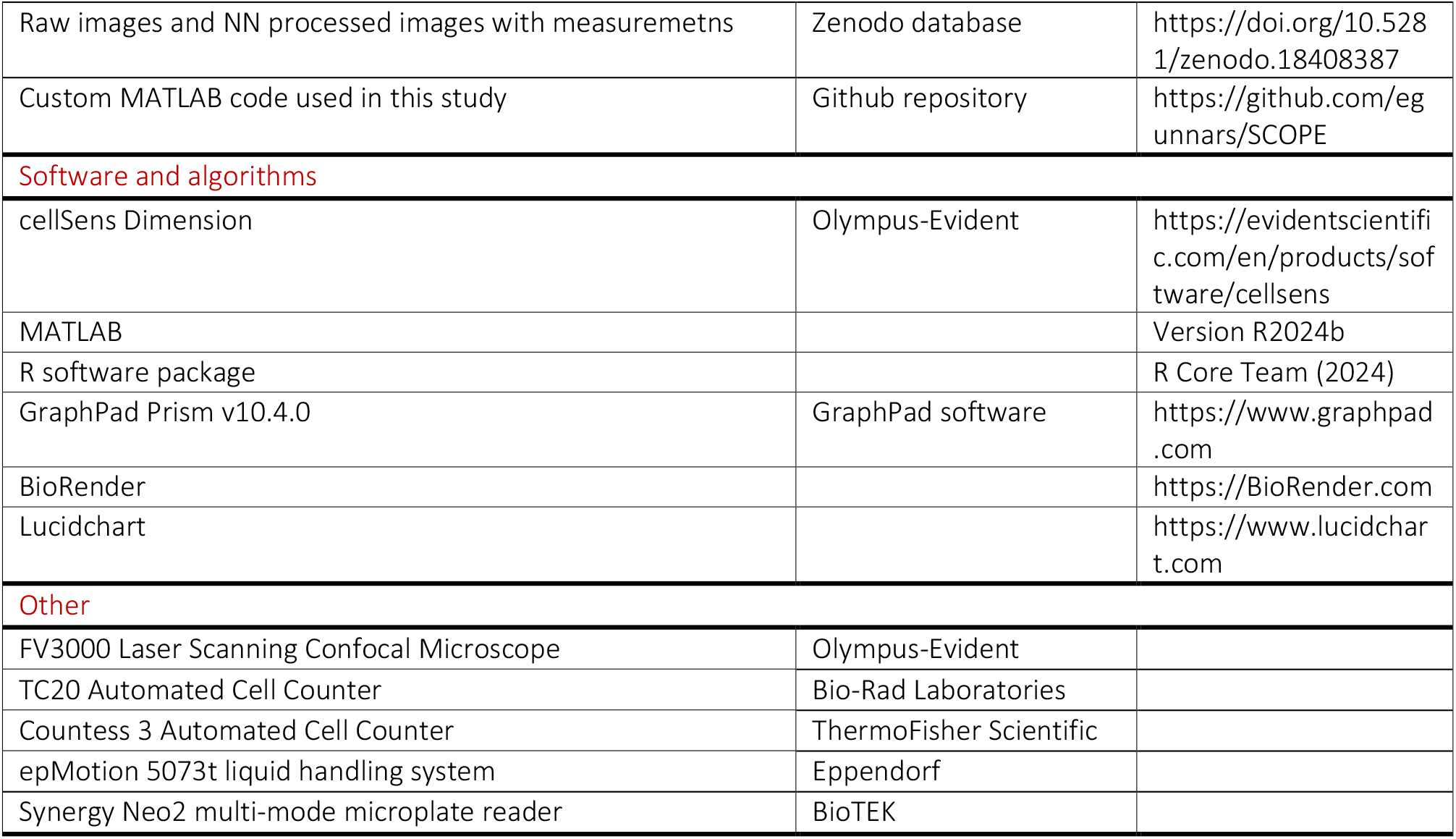

## RESOURCE AVAILABILITY

### Lead contact

Further information and requests for reagents should be directed to and will be fulfilled by the lead correspondance.

### Materials availability

Specific material is available from the lead contact with a completed Materials Transfer Agreement.

### Data and code availability

- All the raw data reported in this paper will be shared by the lead contact upon request.
- All the raw images and NN processed images are available at Zenodo database: (https://doi.org/10.5281/zenodo.18408387).
- The custom code used in this study is available at Github repository: (https://github.com/egunnars/SCOPE).
- Any additional information required to reanalyze the data is available from the lead contact upon request.

## SUPPLEMENTAL TEXT

### Dynamic, multi-modal, imaging-based PDO drug evaluation platform for phenotypic response analysis

We established a standardized multi-modal drug screening workflow including label-free 3D imaging and AI-analysis of patient-derived organoids (PDOs) across multiple timepoints (pre-treatment Day 0, and post-treatment Days 3 and 5) followed by traditional cell viability measurements using the Cell Titer Glo (CTG) assay (**Figure S1**). To analyze the imaging data, a pre-trained neural network (NN) was used with a macro function to automate image processing and data export (**Video S1**). The pre-trained NN detected individual PDOs and classified them as live or dead over time from DMSO-and staurosporine (STA, pan-kinase inhibitor)-treated conditions (**Figure S1B**). PDOs after STA treatment were classified as dead, while most PDOs were classified as live in the DMSO control group. To validate the accuracy of the NN, we compared the live/dead classifications to cell viability readouts with CTG and to vital dye staining with DRAQ7, a far-red fluorescent DNA dye that stains the nuclei in dead cells. CTG showed very low signals in 1 µM STA-treated US PDOs compared to the untreated conditions suggesting that no live cells were present. We also confirmed that the addition of the DRAQ7 dye did not affect the cell viability results (**Figure S1C**). After imaging both brightfield and DRAQ7 channels, PDOs were classified as live or dead by NN processing (**Figure S1D**), and the quantification of DRAQ7-positive PDOs for each condition were compared (**Figure S1E**). US PDOs classified as live in untreated conditions showed minimal DRAQ7 staining (∼6 % positive). Some minor positive DRAQ7 signal may occur in live PDOs due to natural cell turnover. US PDOs classified as dead in untreated conditions also did not show much DRAQ7 positive signal (∼8 % positive) suggesting that these PDOs might be misclassified by the NN due to their small size. Brightfield imaging showed no live PDOs after STA treatment, which is consistent with the CTG results. In addition, approximately 50 % of dead-classified PDOs were stained positively for DRAQ7. This suggests that the NN-based classification of live/dead status correlates well with conventional viability assays. However, complete correlation was challenging due to the scale of viability detection: CTG readouts reflect population-level effects of PDOs, while DRAQ7 stains DNA only in membrane-compromised cells, and its signal can diminish over time depending on the stage of cell death.

### Heterogeneities of each phenotypic class from SCOPE

While the four drug response classes capture high-level variations in phenotypic response, the growth-viability (*GV)* plot also revealed key variations in response strength between drugs of the same phenotypic response class. For example, thiotepa is a cytotoxic drug like osimertinib, displaying early cytotoxicity. However, under thiotepa, there was minor organoid growth on Day 3 (higher growth score than osimertinib) suggesting that the cytotoxic effects of the drug were milder than osimertinib (**Figure S5, B**). There was no significant change in live PDO percent from Day 3 (osimertinib: 19 % mean live, thiotepa: 26 % mean live) to Day 5 (osimertinib: 16 % mean live, thiotepa: 19 % mean live) between thiotepa and osimertinib because most PDOs died immediately after treatment (**Figure S5C**). Clofarabine is classified as a cytostatic plus cytotoxic drug, similar to niraparib. PDO growth was not observed with clofarabine (lower growth score than niraparib) **(Figure S5D, E)**. While niraparib decreased live PDOs (Day 3: 60 %, Day 5: 20 %), clofarabine did not change the percentage of live PDOs from Day 3 (36 % mean live) to Day 5 (31 % mean live), suggesting that cytotoxic effects of clofarabine were stronger and most PDOs had already died on Day 3 (**Figure S5F**). Trifluridine is a late cytotoxic drug like fludarabine. The organoid growth pattern with trifluridine was similar to that of fludarabine, but there was significantly less PDO growth until Day 3 (lower growth score). No further growth was observed on Day 5 because of the late cytotoxicity. Most PDOs were classified as dead on Day 5. (**Figure S5G, H**). Trifluridine showed a large decrease of live PDOs on Day 5 (trifluridine: 28 % mean live, fludarabine: 23 % mean live) compared to Day 3 (trifluridine: 84 % mean live, fludarabine: 97 % mean live) which is characteristic of this class drugs (**Figure S5I**). Drugs in this class require a certain time to display cytotoxic effects, emphasizing the need of multi-timepoint dynamic imaging to capture temporal information of the drug responses. Pazopanib and alpelisib are medium and weak cytostatic drugs, respectively, compared to zanubrutinib **(Figure 3M)**. Most PDOs with these two drugs were live on Day 5, but the sizes of PDOs were significantly smaller compared to DMSO controls. They were not as small as zanubrutinib-treated PDOs though, suggesting that these drugs inhibit PDO growth less than zanubrutinib. Alpelisib was more effective than pazopanib in inhibiting PDO growth because the average PDO size was smaller than pazopanib (**Figure S5J, K**). Both pazopanib and alpelisib did not decrease live PDOs over time (pazopanib: 92 % to 94 %, alpelisib: 88 % to 89 %), suggesting no significant cytotoxic effects (**Figure S5L**).

## METHOD DETAILS

### Generation of colorectal cancer patient organoids

CRC patient tissue was received from USC Norris Comprehensive Cancer Center after surgical resection and processed as previously described ^1^. Patients were consented and tissue samples were collected under an institutional review board (IRB)-approved protocol at the University of Southern California. All samples were de-identified. The tissue was digested enzymatically and dissociated single cells were grown to organoids in Cultrex Reduced Growth Factor Basement Membrane Extract, Type 2 (BME) (R&D Systems). Patient organoids were cultured (37°C, 5% CO_2_) and maintained in 24-well culture plates.

### Organoid seeding and culture process

Organoid-BME mixture was collected with Gentle Cell Dissociation Reagent (StemCell) and continuously mixed on a rocker at 4°C to dissolve the BME for 30-45 min. Organoids were put in digesting solution (1:1 TrypLE and PBS plus 10 µM Y-27632) at 37 °C for 1-5 min and pipetted to dissociate into a single-cell suspension. Once dissociated, Colon Tumor Organoid growth media (Advanced DMEM/F12, 10 % Fetal bovine serum, 1% Penicillin/Streptomycin, 1 % GlutaMAX, 1 % HEPES, 1X B27, 100 ng/ml Noggin, 1X N2 supplement, 10 mM Nicotinamide, 1 mM N-Acetylcysteine, 500 nM A83-01, 50 ng/ml EGF, 10 µM SB202190) was added to the suspension to neutralize the TrypLE. After filtering the solution through 40 µm cell strainer (Corning, Cat# 352340), live cell numbers were counted using either a TC20 (BioRad) or a Countess 3 (Invitrogen) automated cell counter. Mixtures of 1,000 live cells in 10 µl BME and 0.25 mg/ml collagen I were seeded into each well of a 96-well culture plate (Corning, Cat# 3903) manually or using an epMotion liquid handling system (Eppendorf). 200 µl of Colon Tumor Organoid growth media was added to each well to support organoid growth. After culturing for 4 days, Day 0 imaging was conducted before drug treatment. Day 0 images were analyzed to measure organoid average size and number of organoids in each well as QC to determine if the drug treatment experiment could continue.

### Drug management and treatment

FDA-approved oncology drug plates (AODX) were ordered from the National Cancer Institute (NCI). 5FU (Cayman, Cat# 14416), SN38 (Sigma, Cat# H0165), STA (Sigma, Cat# 569396) were ordered separately as powder or liquid and reconstituted with DMSO for further dilutions. Gefitinib (Sigma, Cat# SML1657) and osimertinib (Selleckchem, Cat#S7297) were re-ordered as powder. For the initial screening with the drug plate, 10 mM stock was diluted to make 10 µM drug solutions in the culture media. 0.1% DMSO in media was used as a vehicle control. For the dose-response experiments of selected drugs, drug powders were dissolved in DMSO at maximum soluble concentrations. Drug dilutions were made to treat the different organoid lines with defined concentrations to perform dose response assays. After the Day 0 imaging, media containing drugs were added to each well using the Eppendorf epMotion liquid handling system.

### 3D confocal imaging and image analysis

3D confocal imaging was performed on Day 0, 3 and 5 after drug treatments using Olympus FV3000 laser scanning confocal microscope system. Day 0 imaging was done right before drug treatments and following Day 3 and Day 5 imaging measured the drug effects. Resonant scanner was used for fast scanning with 4 frame average. 512×512 pixel resolution images were captured for multi-z stacks with 640 nm laser illumination in transmitted detector channel for label-free organoid imaging. Each plate was imaged with same plate map and protocol, so the scanning position, the number of z slices and laser intensity were all uniform throughout the experiments. The previously trained NN ^2^ was applied to classify label-free organoid images and the cellSens Tru-AI module with macro (Olympus-Evident) was used to automate the batch processing of organoid imaging data.

### Cell viability assay

After Day 5 imaging, culture media were removed from each well and replaced with 100 µl basal media (Advanced DMEM/F12). 100 µl of CellTiter-Glo (CTG) 3D (Promega) reagent was added to each well, the covered plate was shaken for 5 min and incubated for 25 min in the dark. Relative luminescence units (RLU) values from all wells were read with Synergy Neo2 multi-mode microplate reader (BioTEK). Data was exported as excel spreadsheets and dose-response curves were generated using MATLAB version R2024b.

### SCOPE analysis of PDOs with control drugs

Nine conditions (DMSO vehicle and eight serial dilutions) of SN38 and 5FU were tested in six PDOs (U7, UA, UP, US, UX, V8). Day 0, 3 and 5 images were combined as a time series. Time series images were processed with the trained NN and individual organoids were tracked over time using the cellSens software. Fully tracked organoids with area ≥ 300 µm^2^ at all time points were used as input data for mathematical modeling. Organoids that were classified as dead on Day 3 and live on Day 5 were excluded from the analysis. DRCs and bar graphs with CCTR values were generated using MATLAB version R2024b.

#### [Calculation of PDO growth score]

The growth score was computed based on organoid area changes, but it can equivalently be viewed as describing volume changes. If each organoid is an ellipsoid and the third axis is the geometric mean of the other two axes, its volume is related to its area via *V(t) = 4/(3π^1/2^) × A(t)^3/2^*. If *A*(*t*) follows the Gompertz model, then *V*(*t*) also follows the Gompertz model with initial size *V_0_ = 4/(3π^1/2^) × A_0_ ^3/2^* and growth parameters *a’ =* (3/2) *× a* and *b*^ʹ^ = *b*. Since the ratio *a’/b’* = (3/2) *×* (*a/b*) is a simple linear scaling of the original ratio *a/b*, the normalized growth score remains unchanged if organoid area is replaced by organoid volume.

#### [Growth and viability score]

The Gompertz growth model was used to describe the growth trajectories of individual organoids ^2^. Under this model, the organoid grows initially at exponential rate *a,* and the growth rate decays over time according to an exponential decay parameter *b* ≥ 0. If *A*_0_ is the initial area of an organoid, its area *A*(*t*) at time *t* is given by

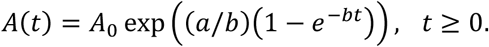

Eventually, the organoid stops growing and reaches a final size *K* = *A*_0_exp(*a/b*) which is the “carrying capacity”. The ratio *a/b* = log(*K/A*_0_) between the growth and decay parameters is a measure of how much an organoid grows in total. Each fully tracked organoid, irrespective of its viability status on Day 5, was fitted with the Gompertz model to obtain estimates of *a* and *b*. For some organoids, the decay parameter *b* was effectively zero (*b* = 10^−3^), in which case the organoid was treated as exponentially growing or decaying with rate parameter *a*. MATLAB version R2024b was used for the parameter fitting. For each patient, each drug and each drug dose, the estimated Gompertz parameters of the fully tracked organoids across three technical replicates were used to compute a growth score. For nonexponential organoids (*b >* 10^−3^), a growth score GS_nonexp_ was computed as the interquartile mean of *a/b* across all individual organoids. Growth scores for DMSO and STA were used to normalize the outcome so that DMSO had a growth score of 1 (drug-free growth) and STA had a growth score of 0 (complete growth inhibition). The normalization was done on a plate-by-plate basis. For exponential organoids (*b* = 10^−3^), a growth score GS<u>_exp_</u> was computed in the same way, with the exponential growth rate *a* in place of the ratio *a/b*, and then normalized using DMSO and STA. A final growth score was computed as follows:

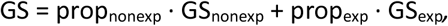

where prop_nonexp_ and prop_exp_ were the proportions of nonexponential and exponential organoids, respectively, across fully tracked organoids for the condition in question. For each patient, each drug and each drug dose, a 95 % confidence interval for the growth score was computed using the bootstrap method.

A viability score was calculated based on the live and dead classifications by the NN. Among the fully tracked individual organoids, we calculated the proportion of organoids that kept live status between Day 3 and Day 5 (live-live organoids) and used the value to represent organoid viability. The viability score was normalized using live-live percentages under DMSO and STA, so that DMSO would have a viability score of 1 and STA a viability score of 0.

#### [2D DRC and CCTR]

The growth score and the proportion of live-live tracked organoids (viability score) were plotted as a function of dose to produce two distinct dose response curves (DRCs), separating the effect of the drug on organoid growth and viability. The upper horizontal asymptote of each DRC was assumed to be 1. Each DRC was used to define an IC_50_ dose, the dose achieving a growth score of 0.5 and a live-live percentage of 50 %, respectively, relative to DMSO and STA. The logarithmic difference between the two IC_50_ doses was defined as the Cytostatic-cytotoxic transition range (CCTR):

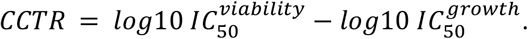

### Single dose screening with phenotypic classifications

#### [Endpoint screening]

Initial screening was performed using Day 5 CTG and NN analysis results. Prior to the single-dose screen, assays were tested for well position effects with the two metrics and the liquid handler was optimized, assuring that percent coefficient of variation (%CV) was below 20 % for PDO under control conditions. Quality assurance was completed using a modified Z-factor, derived from the work of Zhang & colleagues, and % CV ^3^. Single dose (10 µM) of 166 drug compounds were tested in UP PDOs. The experiments were performed in multiple 96 well plates with three technical replicates and each plates included DMSO (Vehicle), STA (5 µM), SN38 (0.01 µM) and 5FU (10 µM) control drugs to compare and define the dynamic response ranges. Test drugs with a negative Z-factor and at least one technical replicate above the percent viability of DMSO minus three times the standard deviation were removed from the dataset or repeated. Plates with a % CV of the control wells greater than 20 % were repeated. CTG and image-analysis test-drug results were normalized to both a positive (STA) and a negative (DMSO) control to calculate percent control viability. For image-analysis data, the percent control viability was calculated from the total area of live organoids per-well with Day 5 images. Test drugs with percent viability values below a standard cutoff of the negative (100 %) DMSO control minus three standard deviations in both CTG and image-analysis assays were considered hit drugs. CTG and image quality assurance and statistical analysis were performed using R version 4.4.0 in RStudio ^4^.

#### [SCOPE analysis]

Growth scores and 95% confidence intervals for each of the 166 FDA-approved anti-cancer compounds applied to the UP PDO line were computed as described above. Time series tracking data of organoid viability was combined with the growth scores to define a new summary statistic of dynamic drug response, where the response of each drug was summarized using a 3D vector (*x, y, z*). The *x*-coordinate was the proportion of tracked organoids that were live on Day 3 and Day 5 (live-live), the *y*-coordinate was the proportion of tracked organoids live on Day 3 and dead on Day 5 (live-dead), and the *z*-coordinate was the growth score computed as above. This enabled visualization of the drug response using a 3D *GV (growth-viability)* plot. The *GV* plot was generated using MATLAB version R2024b. The GV plot was used to define four phenotypic classes of drug response: cytotoxic, late cytotoxic, cytostatic, and cytotoxic + cytostatic. The definitions of the classes are as follows, where LL denotes the percentage of live-live tracked organoids, LD denotes the percentage of live-dead tracked organoids, and DD denotes the percentage of dead-dead organoids (dead on Day 3 and dead on Day 5):

- Cytotoxic: DD ≥ 0.7.
- Late cytotoxic: LD ≥ 0.15 and LD ≥ (4/5) − (5/6)•LL.
- Cytostatic: LL ≥ 0.75 and LD ≤ 0.15.
- Cytostatic + cytotoxic: All drugs not falling into any of the previous three categories.

The cytostatic category was further subdivided as follows:

- Weak to no effect: Growth score above 0.7.
- Medium effect: Growth score between 0.3 and 0.7.
- Strong effect: Growth score below 0.3.

Typical example drugs from each class were selected to represent the organoid growth inhibition and cytotoxic effects. NN-processed time series images, organoid area measurements and the percentages of live/dead organoids between Day 3 and Day 5 were used to compare the characteristics of phenotypic drug responses of each class. SCOPE phenotypic classifications of hit drugs (124 drugs) were compared to manual review results by organoid experts to validate the effects of each class drugs. Using the NCI drug plate information including target molecules and biological pathways, drugs in the four classes were grouped into specific categories.

### SCOPE analysis of EGFR inhibitors

Gefitinib and osimertinib were tested in six PDOs (U7, UA, UP, US, UX, V8) with eight different drug concentrations. SCOPE analysis was performed same as for the control drugs. 2D DRCs were generated to calculate the IC_50_ and CCTR values. 3D *GV* plots were generated with multiple doses of gefitinib and osimertinib to track the changes in phenotypic response. Time series tracking data of organoid viability was used to classify each condition into one of the four phenotypic classes as described above. Fitted growth score DRCs were used to subclassify cytostatic drugs into strong, medium and low to no effect.

## Supplemental information

**Figure S1:**
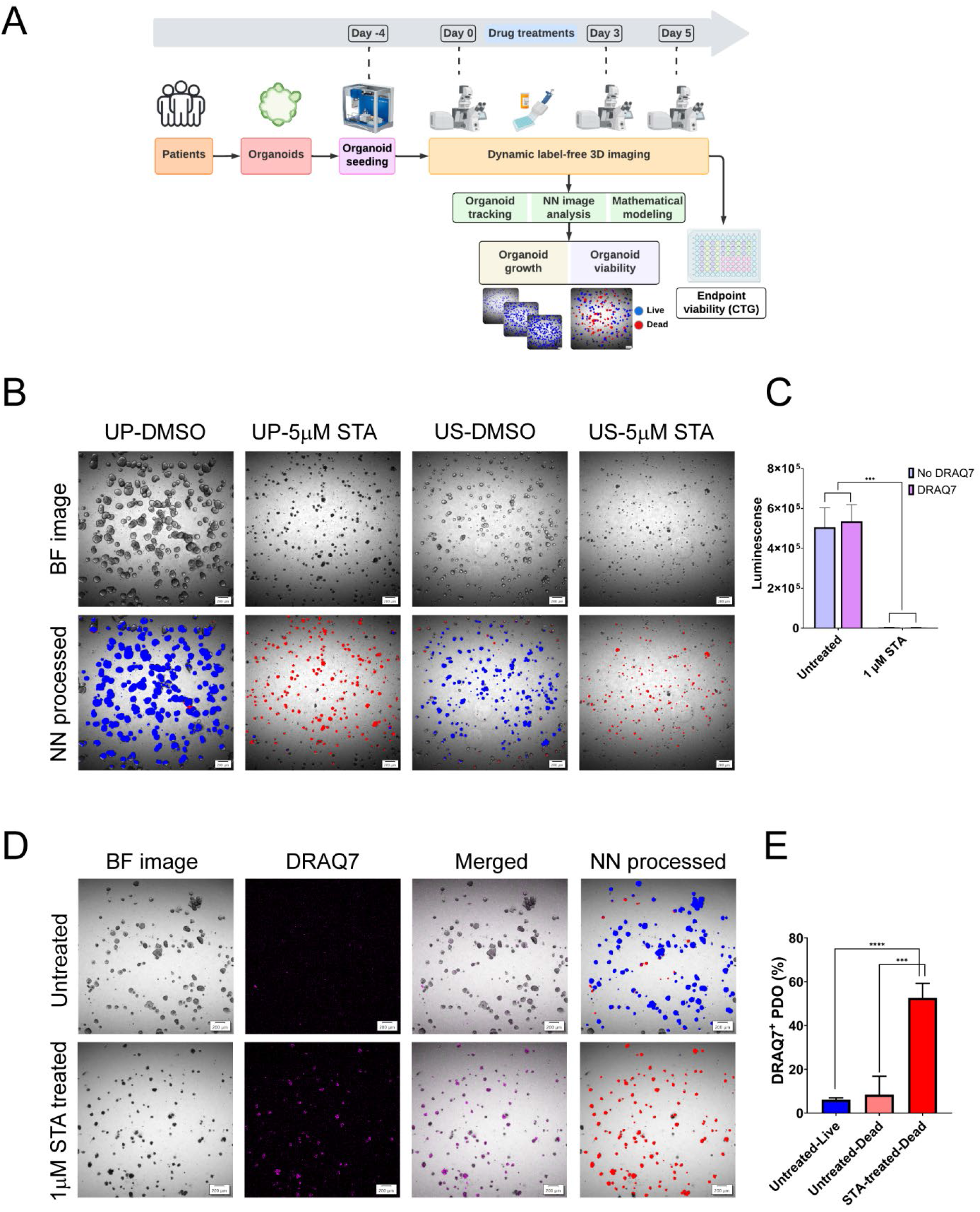
Dynamic, multi-modal imaging-based PDO drug screening. **A.** The workflow includes optimized PDO handling, multi-timepoint label-free 3D imaging, NN-based image analysis, and CTG cell viability assay. Organoid growth and viability information were used to evaluate drug effects in PDOs. Created with Lucidchart software (https://www.lucidchart.com) **B.** NN processing results of UP and US PDO brightfield (BF) images with DMSO control and STA-treated conditions. **C.** CTG results of untreated and 1µM STA-treated US PDOs with or without DRAQ7 dye. (CTG signals, No DRAQ7-Untreated: 505,617.8 ± 96,612.47, No DRAQ7-1µM STA: 3,665.8 ± 1,373.97, n = 5, *\*\*\*p=0.0003*, DRAQ7-Untreated: 535,743.2 ± 83,017.99, DRAQ7-1µM STA: 2,986.4 ± 1,480.74, n = 5, *\*\*\*p=0.0001*). Mean ± SD. Paired t-test. **D.** BF and DRAQ7-stained images for untreated and STA-treated US PDOs with NN live/dead classifications. **E.** Quantification of DRAQ7-positive PDOs in NN-classified live/dead PDOs. (Untreated-Live: 6.10 % ± 0.80, Untreated-Dead: 8.39 % ± 8.41, STA-treated-Dead: 52.65 % ± 6.59, n = 4, Untreated-Live vs. STA-treated-Dead, *\*\*\*\*p<0.0001*, Untreated-Dead vs. STA-treated-Dead, *\*\*\*p=0.0007*, Unpaired t-test. Scale bars, 200 µm.

**Figure S2:**
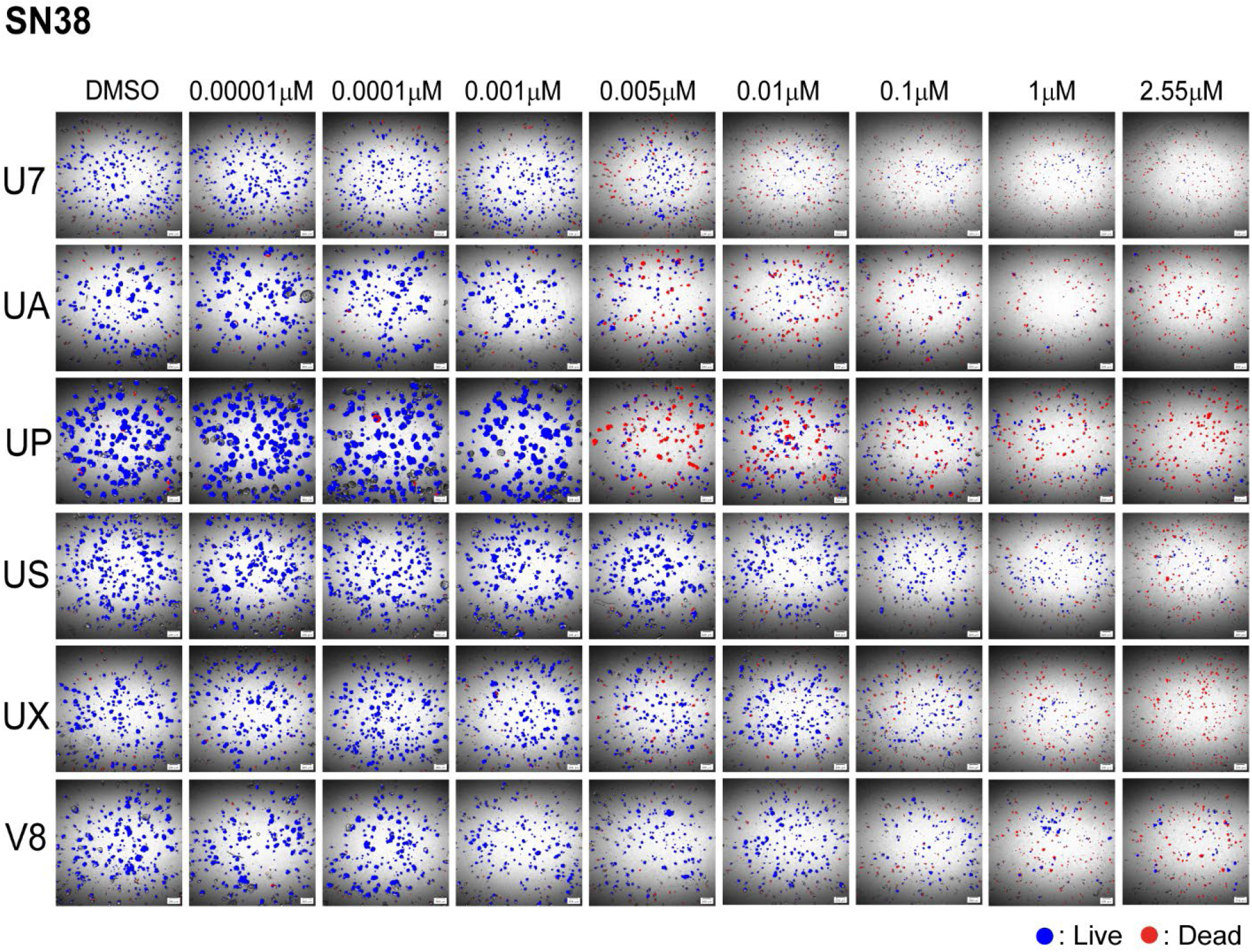
SN38 dose response in six PDOs. Day 5 NN-processed images of six PDOs treated with eight different doses of SN38 plus DMSO as vehicle no-treatment control. Live (Blue) and dead (Red) organoids were classified by the pre-trained NN. Scale bars, 200 µm.

**Figure S3:**
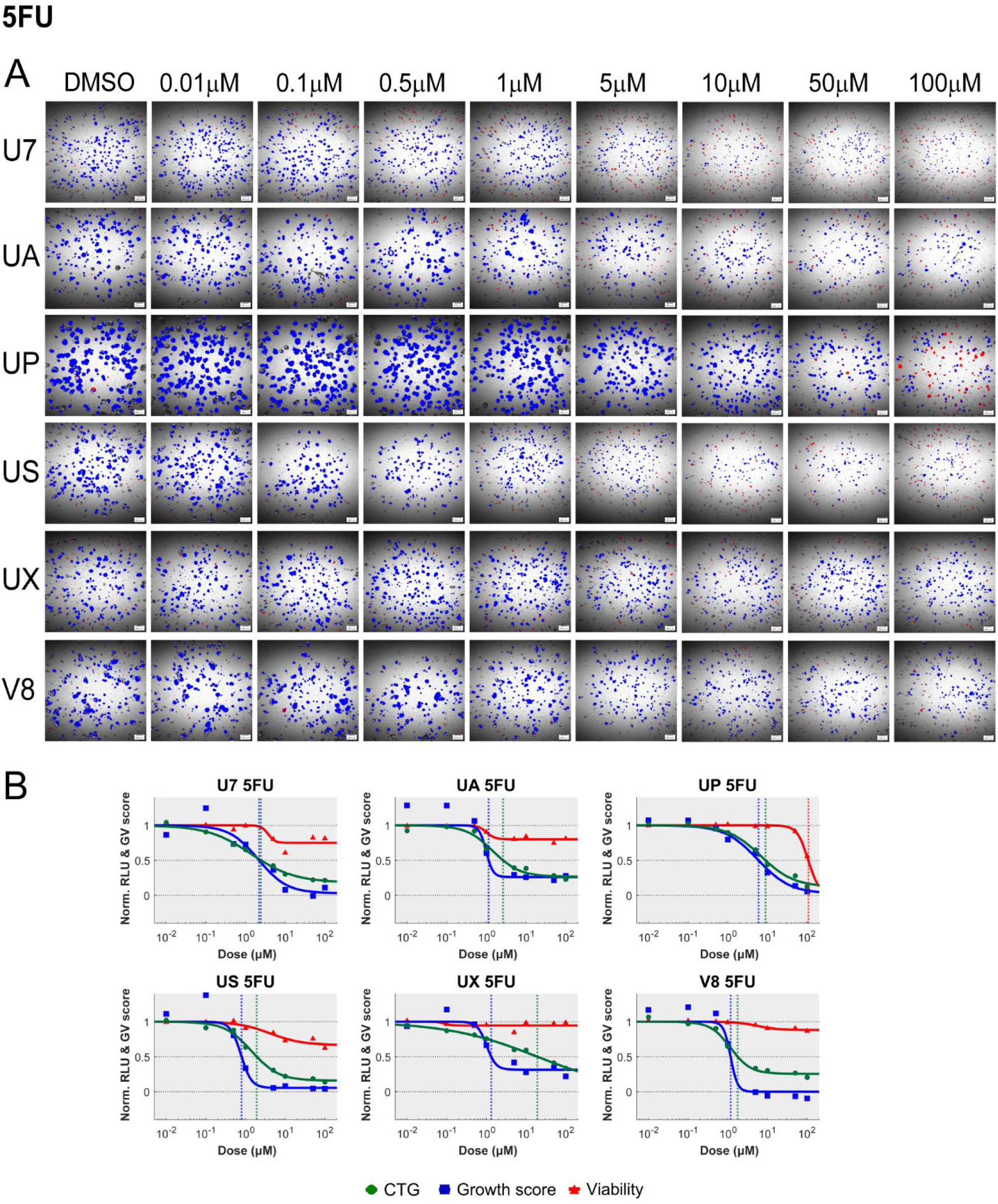
5FU dose response in six PDOs. **A.** Day 5 NN-processed images of six PDOs treated with eight different doses of 5FU plus DMSO as vehicle no-treatment control. Live (Blue) and dead (Red) organoids were classified by the pre-trained NN. Scale bars, 200 µm. **B.** 5FU dose response curves of six PDOs based on CTG (Green), growth (Blue) and viability (Red) scores. Vertical blue and green dotted lines indicate IC_50_-growth and IC_50_-CTG values. Red dotted lines for IC_50_-viability are omitted where 50% viability is not achieved.

**Figure S4:**
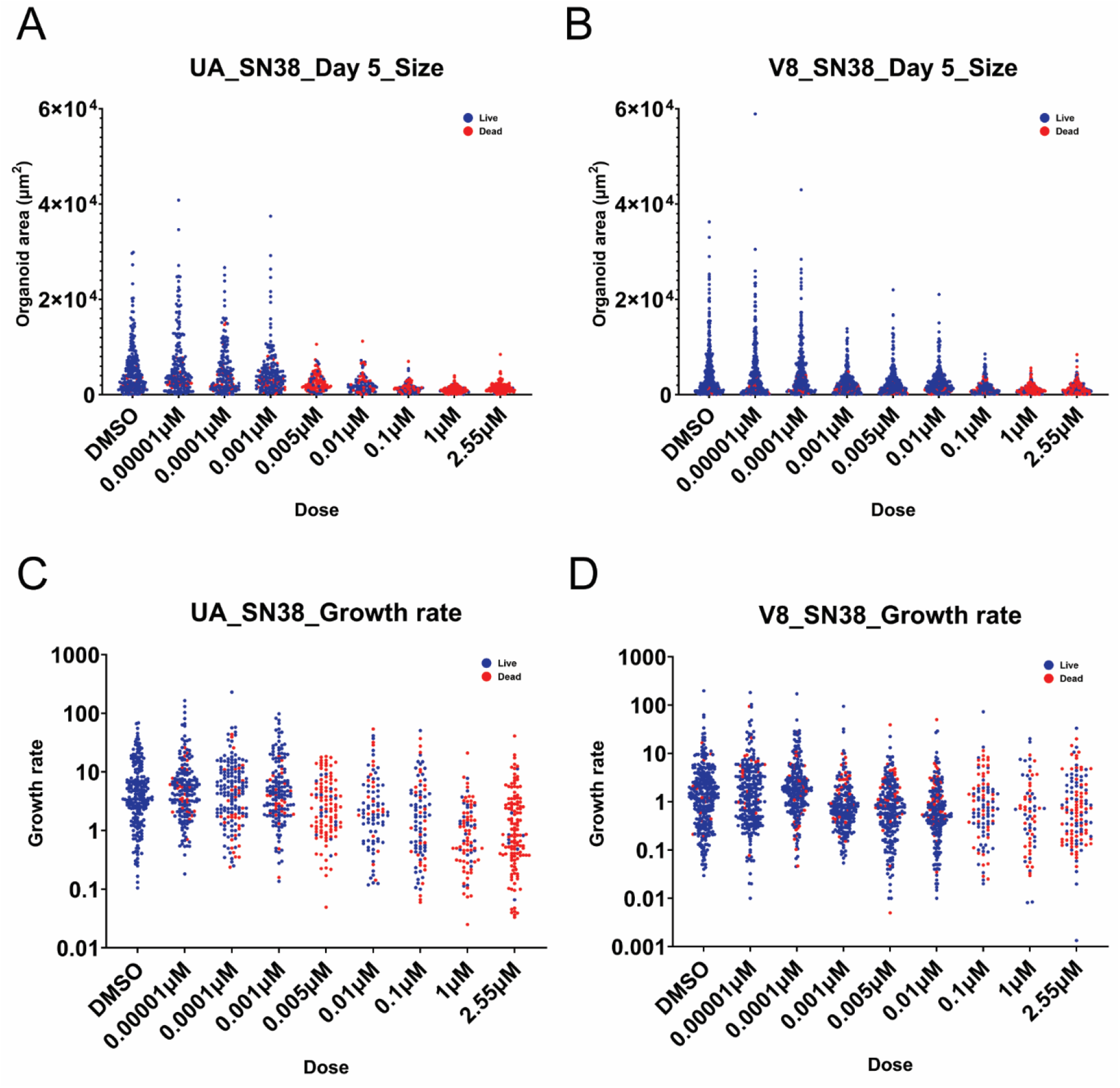
SN38 dose response of individual organoids in UA and V8 PDOs. A, **B.** Size distributions of individually tracked organoids over multiple doses of SN38 in UA and V8 PDOs. **C, D.** Growth rate measurements of individual organoids in UA and V8 PDOs treated with SN38. Blue: live PDOs, Red: dead PDOs. Each dot represents a single individual organoid. Growth rate = log_10_ [Day 5 area-Day 0 area) / Day 0 area].

**Figure S5:**
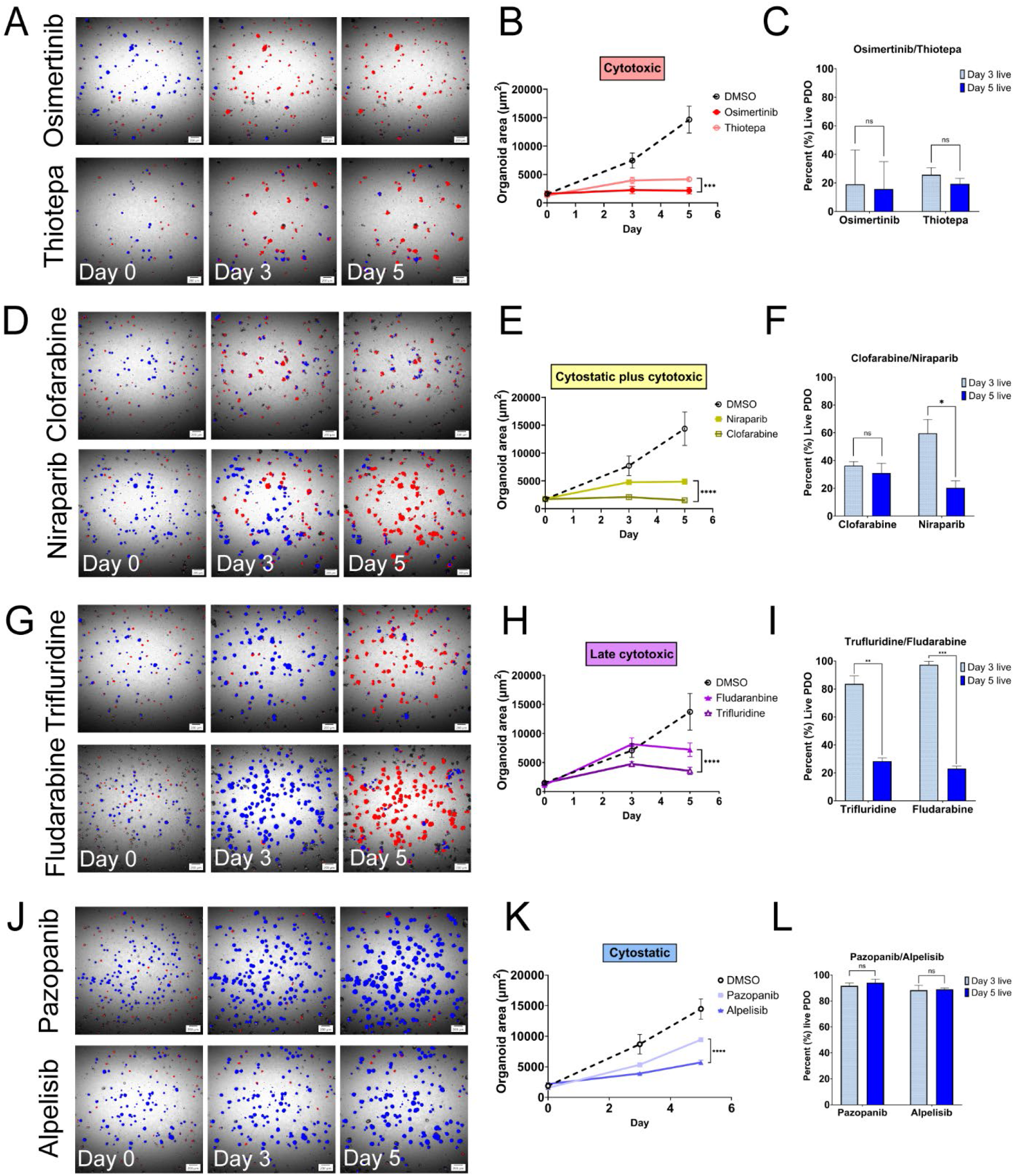
Intra-class heterogeneities of phenotypic classifications. **A.** Representative NN-processed images of osimertinib-and thiotepa-treated PDOs. **B.** PDO area changes over time with osimertinib and thiotepa treatments. (DMSO Day 5: 14,652.42 µm^2^ ± 2,364.65 µm^2^, Osimertinib Day 5: 2,157.28 µm^2^ ± 591.64 µm^2^, Thiotepa Day 5: 4,150.26 µm^2^ ± 261.49 µm^2^, n = 3, *\*\*\*p=0.0003*). Mean ± SD. Two-way ANOVA test. **C.** Percent changes of live PDOs with osimertinib and thiotepa between Day 3 and 5. (Osimertinib Day 3: 19 % ± 24.02 %, Osimertinib Day 5: 15.67 % ± 19.35 %, n = 3, *^ns^p=0.5085*, Thiotepa Day 3: 25.67 % ± 5.03 %, Thiotepa Day 5: 19.33 % ± 3.79 %, n = 3, *^ns^p=0.0890*). Mean ± SD. Paired t-test. **D.** Representative NN-processed images of clofarabine and niraparib-treated PDOs. **E.** PDO area changes over time with clofarabine and niraparib treatments. (DMSO Day 5: 14,354.6 µm^2^ ± 2,998.30 µm^2^, Clofarabine Day 5: 1,521.21 µm^2^ ± 117.65 µm^2^, Niraparib Day 5: 4,858.67 µm^2^ ± 491.18 µm^2^, n = 3, *\*\*\*\*p<0.0001*). Mean ± SD. Two-way ANOVA test. **F.** Percent changes of live PDOs with clofarabine and niraparib between Day 3 and 5. (Clofarabine Day 3: 36.33 % ± 2.89 %, Clofarabine Day 5: 31 % ± 7 %, n = 3, *^ns^p=0.4158*, Niraparib Day 3: 59.67 % ± 10.02 %, Niraparib Day 5: 20.33 % ± 5.03 %, n = 3, *\*p=0.0197*). Mean ± SD. Paired t-test. **G.** Representative NN-processed images of trifluridine-and fludarabine-treated PDOs. **H.** PDO area changes over time with trifluridine and fludarabine treatments. (DMSO Day 5: 13,709.69 µm^2^ ± 3,154.03 µm^2^, Trifluridine Day 5: 3,568.45 µm^2^ ± 639.62 µm^2^, Fludarabine Day 5: 7,210.79 µm^2^ ± 1,171.45 µm^2^, n = 3, *\*\*\*\*p<0.0001*). Mean ± SD. Two-way ANOVA test. **I.** Percent changes of live PDOs with trifluridine and fludarabine between Day 3 and 5. (Trifluridine Day 3: 83.67 % ± 5.69 %, Trifluridine Day 5: 28.33 % ± 2.31 %, n = 3, *\*\*p=0.0061*, Fludarabine Day 3: 97.33 % ± 2.52 %, Fludarabine Day 5: 23 % ± 1.73 %, n = 3, *\*\*\*\*p<0.0001*). Mean ± SD. Paired t-test. **J.** Representative NN-processed images of pazopanib-and alpelisib-treated PDOs. **K.** PDO area changes over time with pazopanib and alpelisib treatments. (DMSO Day 5: 14,472.74 µm^2^ ± 1,657.88 µm^2^, Pazopanib Day 5: 9,436.51 µm^2^ ± 119.67 µm^2^, Alpelisib Day 5: 5,712.56 µm^2^ ± 361.22, µm^2^ n = 3, *\*\*\*\*p<0.0001*). Mean ± SD. Two-way ANOVA test. **l.** Percent changes of live PDOs with pazopanib and alpelisib between Day 3 and 5. (Pazopanib Day 3: 91.53 % ± 2.33 %, Pazopanib Day 5: 94 % ± 2.82 %, n = 3, *^ns^p=0.3785*, Alpelisib Day 3: 88.17 % ± 3.90 %, Alpelisib Day 5: 88.87 % ± 1.00 %, n = 3, *^ns^p=0.7799*). Mean ± SD. Paired t-test. Scale bars, 200 µm.

**Figure S6:**
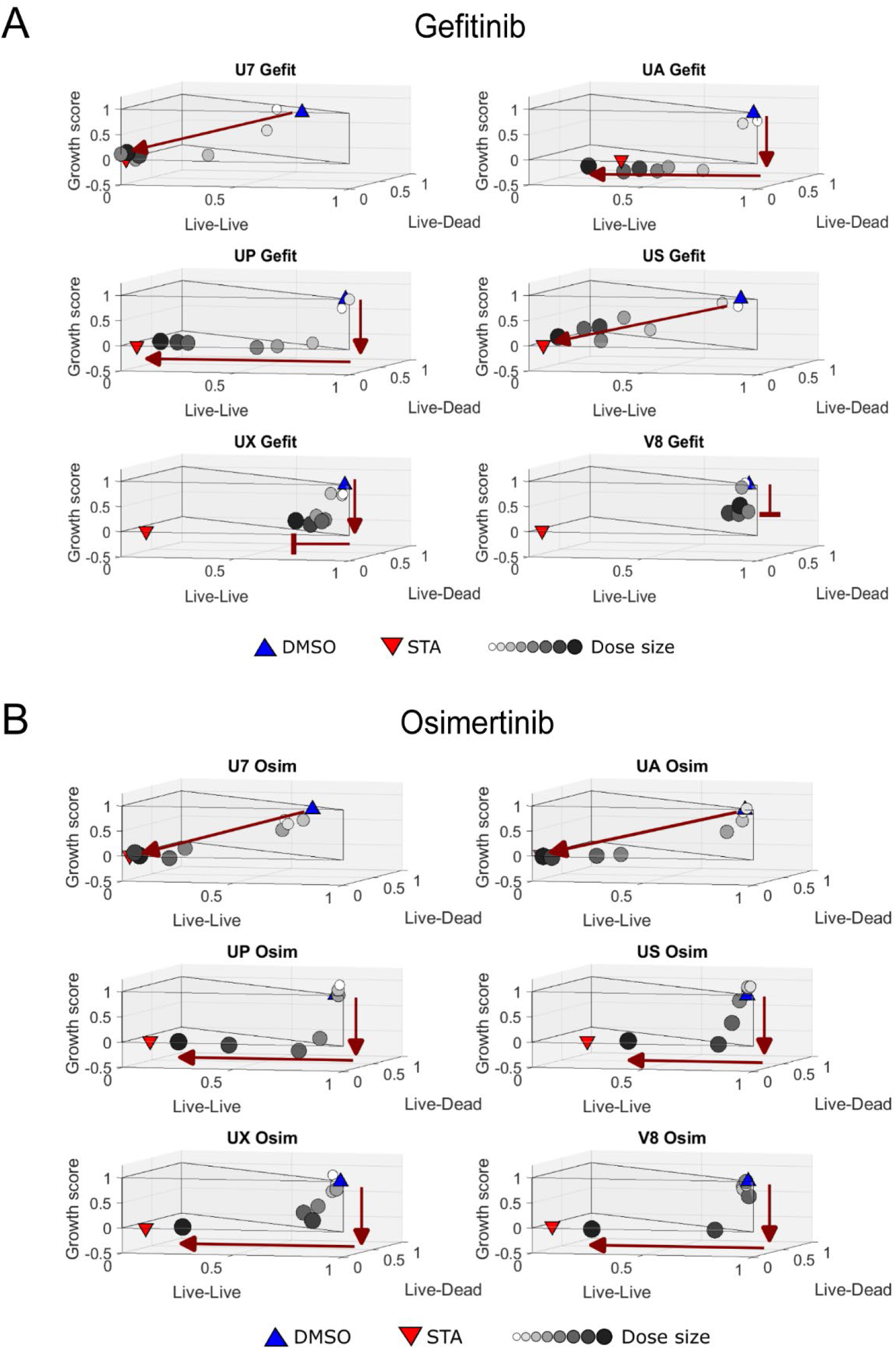
Dose-dependent phenotypic changes of six PDOs with gefitinib and osimertinib. 3D *GV* plots of U7, UA, UP, US, UX and V8 PDOs with gefitinib **(A)** and osimertinib **(B)** over multiple doses. Dark red arrows show the dose-dependent *GV* score trajectories. Light gray to black circles visualize each dose data point from low to high concentrations.

**Table S1:**
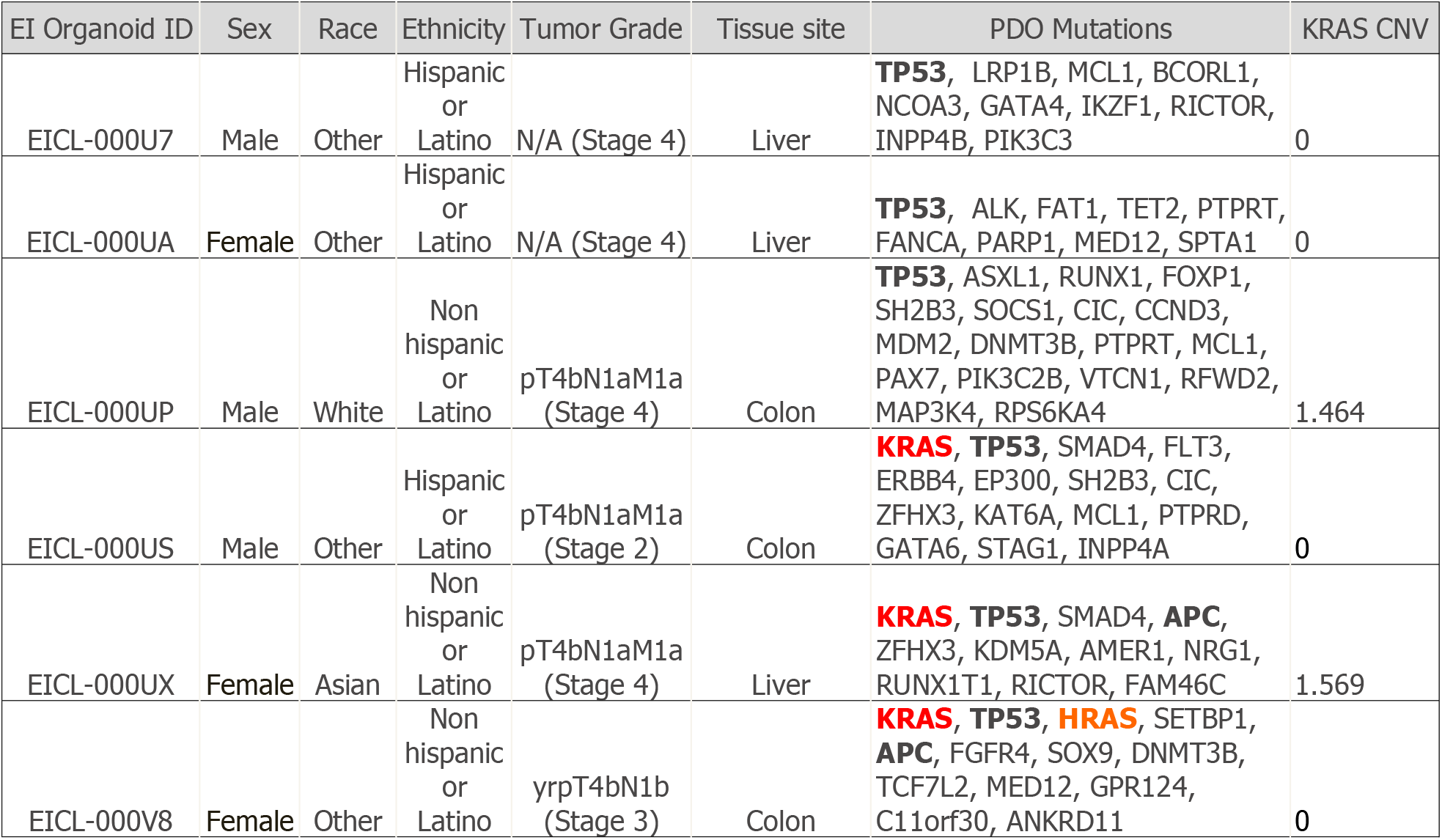
Patient-organoid information and gene mutations.

**Data S1: IC_50_-growth, IC_50_-viability and IC_50_-CTG of SN38, 5FU, gefitinib and osimertinib with six PDOs**

Data S1
| SN38 |  |  |  |
| --- | --- | --- | --- |
|  | IC50-growth | IC50-viability | IC50-CTG |
| U7 | 0.0047 | 0.0409 | 0.0040 |
| UA | 0.0058 | 0.0114 | 0.0049 |
| UP | 0.0025 | 0.0115 | 0.0025 |
| US | 0.0573 | 1.5448 | 0.0028 |
| UX | 0.0057 | 0.1607 | 0.0080 |
| V8 | 0.0007 | 0.3240 | 0.0013 |
| 5FU |  |  |  |
|  | IC50-growth | IC50-viability | IC50-CTG |
| U7 | 2.1949 | ≥ 100 | 2.4528 |
| UA | 1.1407 | ≥ 100 | 2.6171 |
| UP | 5.9430 | ≥ 100 | 8.9119 |
| US | 0.7936 | ≥ 100 | 1.9138 |
| UX | 1.3212 | ≥ 100 | 19.2388 |
| V8 | 1.1759 | ≥ 100 | 1.7743 |
| Gefitinib |  |  |  |
|  | IC50-growth | IC50-viability | IC50-CTG |
| U7 | 0.1105 | 0.2707 | 0.2098 |
| UA | 0.1349 | 0.6525 | 0.1927 |
| UP | 0.3201 | 3.0906 | 0.3733 |
| US | 0.3844 | 0.5142 | ≥ 100 |
| UX | 1.0885 | ≥ 100 | 0.6310 |
| V8 | 2.9519 | ≥ 100 | 5.7640 |
| Osimertinib |  |  |  |
|  | IC50-growth | IC50-viability | IC50-CTG |
| U7 | 0.0815 | 0.2574 | 0.1198 |
| UA | 0.0932 | 0.3734 | 0.0991 |
| UP | 0.3283 | 3.1866 | 0.1869 |
| US | 0.9623 | 7.3871 | 0.5085 |
| UX | 0.4153 | 7.2108 | 0.3667 |
| V8 | 1.3281 | 7.0184 | 1.3413 |

**Data S2: UP PDOs growth scores and viability percentages with screened drugs**

Data S2. UP PDO_Growth scores and viability percentages.xlsx

**Data S3: Phenotypic classifications (SCOPE vs. Manual) and target pathways**

Data S3. Phenotypic classifications_manual_vs_modeling.xlsx

**Video S1: NN processing of time series images and PDO tracking.** The video shows image preprocessing steps, PDO detection and live/dead classifications by NN and PDO tracking of three different timepoints. UP PDOs treated with niraparib (10 µM). EFI: Extended Focal Imaging. Scale bars, 200 µm.

Movie S1.avi

## Notes

### Competing Interest Statement

The authors have declared no competing interest.

